# Cell cycle checkpoint activity in the malaria parasite *Plasmodium falciparum*

**DOI:** 10.1101/2025.05.08.652832

**Authors:** Monique K. Johnson, Juliana Naldoni, William H. Lewis, Ross F. Waller, Catherine J. Merrick

**Affiliations:** Department of Pathology, University of Cambridge, Tennis Court Road, Cambridge, CB2 1QP, UK; Department of Biochemistry, University of Cambridge, Tennis Court Road, Cambridge, CB2 1QW, UK

**Author notes:** Address correspondence to: Department of Pathology, Cambridge University, Tennis Court Road, Cambridge, CB2 1QP, UK.

**Keywords:** Malaria, *Plasmodium*, cell cycle, checkpoint, PI3K

## Abstract

*Plasmodium spp*. have different modes of cell division from most eukaryotes. Little is known about how these are controlled and cell-cycle checkpoints are particularly poorly characterised. However, parasites can arrest their cell cycle when treated with the frontline antimalarial drug artemisinin, and artemisinin-resistant parasites can modulate their cell-cycle progression, so it is important to understand these aspects of *Plasmodium* biology. Here, we show that *P. falciparum* displays hallmarks of an intra-S-phase checkpoint when exposed to DNA damage, including acute reduction of DNA replication and phosphorylation of a putative damage-marker histone. Compounds that inhibit human checkpoint kinases can inhibit this arrest of DNA replication, and synergise with DNA damage in parasite killing. This suggests the existence of checkpoint kinase activity in *P. falciparum*, yet these kinases have no clear homologues in *Plasmodium* genomes. Their closest homologues are the phosphatidylinositol lipid kinases. We hypothesise that phosphatidylinositol 3-kinase – which is reportedly up-regulated in artemisinin-resistant parasites – may moonlight in this role, and we characterise this essential kinase for the first time via expansion microscopy. Finally, we show that the cryptic checkpoint-kinase activity may also regulate the ring-stage survival phenotype after artemisinin damage, which resembles a G1/S checkpoint. Hence we suggest that checkpoint kinase inhibitors are candidates for synergy with artemisinin.

**Importance:** Malaria parasites infect red blood cells, wherein they replicate to produce many new parasites. This is unusual because most cells replicate simply by copying their genome and splitting in half (called binary fission) but malaria parasites make ∼20 genome copies, then partition them simultaneously into 20 new cells (called schizogony). Here we studied how schizogony is controlled: in particular, are there ‘checkpoints’? Checkpoints pause the cycle for repair if the genome is damaged. We found that DNA damage *did* cause checkpoint hallmarks, yet key proteins that enforce this in other cells are absent in malaria parasites. Furthermore, this checkpoint activity may be involved in the response to an antimalarial drug, when parasites pause their cycle before active replication begins. This implies that inhibiting the checkpoint could exacerbate parasite killing by such drugs. Cancer therapies often work like this – by damaging DNA and also preventing the cancer cells from repairing it.

## Introduction

*Plasmodium falciparum* is the most important cause of human malaria, giving rise to over half a million deaths and hundreds of millions of clinical cases per year (1). Besides being a major human pathogen, it is also a highly unusual protozoan organism, distantly related to humans and other model organisms, with distinctive features of cell and molecular biology. One of these is a non-canonical cell cycle. Instead of dividing by binary fission, as in most model organisms, *Plasmodium* parasites divide primarily by schizogony. After invading a host erythrocyte, they execute multiple, asynchronous rounds of genome replication and nuclear division, generating multiple nuclei within the same cytoplasm. This is followed by a relatively synchronous mass cytokinesis event, partitioning ∼20 nuclei and organelles into individual merozoites.

The complex events of schizogony are now being dissected in molecular and cellular detail (2-5) but fundamental questions remain about how the process is regulated. Schizogony is not easily mapped onto the cell cycle phases of G1, S, G2 and M. After invading a host cell, the parasite undergoes a pre-replicative growth phase broadly analogous to G1. The haploid genome is then repeatedly replicated in a continuous S-phase without once-and-only-once regulation, generating many new haploid nuclei. Different nuclei cease to replicate at different times, and divide at different times, so there is no well-defined G2 and no single M-phase (3).

*Plasmodium* replication is not controlled by conventional cyclin-dependent kinase activity (6), nor is it apparently governed by conventional cell cycle checkpoints, which usually operate at the G1/S transition, within S-phase, and at G2/M. Respectively, these checkpoints delay the start of S-phase, arrest replication if DNA damage or replication stress is detected, and prevent mitosis if a genome is incompletely replicated (7). The *Plasmodium* genus, however, encodes no clear homologues of the checkpoint kinases conserved in most eukaryotes (animals, fungi and plants) e.g. *S. cerevisiae* Mec1 and Rad53 or human ATM/ATR and Chk1/Chk2 (6). (By contrast, some parasites in the apicomplexan phylum, like *Toxoplasma*, do retain homologues of the ‘master regulators’ Mec1/ATM/ATR, which are phosphoinositide-3-kinase-like kinases (PIKKs), and checkpoints are better characterised in *Toxoplasma* (8-10)).

Nevertheless, it seems unlikely that *Plasmodium* is entirely without checkpoints. Eukaryotic cells (not only in metazoans, which must guard against cancer, but also protozoans) generally do pause their cell cycles to ensure effective DNA repair. *Plasmodium* encodes most of the known DNA repair pathways (6, 11) and generates relatively low rates of SNPs, indels and translocations during normal replication (12).

There is some evidence for a G1/S-like checkpoint: pre-replicative rings can undergo ‘temporary growth arrest (TGA)’ (13) in response to nutritional (14), temperature (15) or antimalarial-drug (16, 17) stresses. TGA is particularly important under stress caused by the antimalarial drug artemisinin, because it can contribute to treatment failure. However, there is no molecular evidence as-yet that conventional checkpoint effectors are involved. A G1/S checkpoint usually involves the cyclin/CDK complexes that drive S-phase entry, influenced positively by growth factors or negatively by damage responsive kinases like PIKKs. TGA in *Plasmodium*, by contrast, is defined only as a ‘non-specific stress-response survival mechanism’ (13): induced by multiple stresses, possibly via multiple molecular pathways.

An intra-S-phase checkpoint should arrest S-phase specifically after DNA damage or replication stress. In *Plasmodium*, the transcriptional cascade that drives the cell cycle can slow down after DNA damage, with DNA repair proteins and chromatin modifications being upregulated (18), but the signals and transducers for any potential checkpoint are unknown. *Plasmodium* generally lacks validated markers for DNA damage and for direct responses to it. Indeed, the multi-nucleate, asynchronous nature of schizogony raises questions about how an intra-S-phase checkpoint should or could be enforced. Replication itself is apparently regulated per-nucleus, probably via individual accumulations of PCNA, together with non-canonical, chromatin-bound CDK-related kinases like CRK4 (19). Nevertheless, diffusible checkpoint kinases, potentially affecting the whole cell, may not be incompatible with schizogony – at least in response to exogenous DNA damage.

Finally, a G2/M checkpoint may be entirely absent because some *P. falciparum* mutants perform grossly aberrant genome segregation but still undergo cytokinesis (20). In fact, even in non-mutants, certain phases of the lifecycle produce a high rate of zoids – this occurs particularly gametogenesis, wherein cytokinesis proceeds even if genome replication fails entirely (21, 22).

Here, we set out to investigate checkpoint responses to DNA damage in *Plasmodium*, using newly developed molecular and cellular tools. We show that the hallmarks of an S-phase checkpoint do occur in *P. falciparum*, despite the apparent absence of canonical PIKKs.

## Results

### Evidence for DNA-damage-responsive checkpoint activity in *P. falciparum*

We used two complementary tools to investigate the response to DNA damage during S-phase in *P. falciparum*, focussing at the levels of DNA and protein. Firstly, we labelled nascent DNA in replicating parasites with a quantifiable modified nucleotide. This tested whether DNA replication would acutely arrest after damage: a hallmark of the intra-S-phase checkpoint (23, 24). Secondly, we measured phosphorylation of histone H2A, which was recently identified as being analogous to histone H2AX phosphorylation in mammalian cells (25). Canonically, H2AX is phosphorylated at chromatin sites of DNA damage (26, 27). H2AX is absent in *Plasmodium*, but the non-variant histone H2A can nevertheless be phosphorylated (25). Importantly, this indicates an *in trans*, kinase-driven, response that is genuinely ‘checkpoint-like’.

To induce DNA damage, we used the well-characterised alkylating agent methyl methane sulphonate (MMS), which alkylates DNA bases, particularly guanine and adenine (28). In mammalian cells, it induces an intra-S-phase checkpoint (23). In *P. falciparum* parasites, it leads to DNA breakage, as measured by comet assays (29) and concomitantly delays the cell cycle (18), but its effect as a checkpoint inducer has not been explicitly addressed. In the *P. falciparum* 3D7 strain, MMS had a 48h IC_50_ of ∼96µM (Fig S1A). Having established this, we performed highly acute assays over just 1-2h, using 10x or 20x IC_50_, to minimise non-specific or secondary effects. We confirmed that parasites could substantially recover from this level of MMS exposure within one cell cycle, although their maturation was delayed, as reported previously (18), and there was also a reduction in merozoite numbers in mature schizonts (Fig S2).

DNA replication was acutely reduced in the 30 minutes following a 30-minute exposure to MMS at 10x or 20x IC_50_. Some reduction occurred throughout S-phase (∼28-36 hours post invasion, hpi) but it was most pronounced when DNA was damaged in early trophozoites at ∼32hpi (Fig 1A). Within a similar timeframe, histone H2A phosphorylation was moderately elevated (Fig 1B). We recently reported that several DNA damaging agents, including MMS, could cause H2A phosphorylation (30), but that the upregulation in trophozoite stages was modest – usually less than 2-fold over background levels – making it difficult to measure consistently. This was equally true when MMS was used to induce H2A phosphorylation here (Fig 1B). (Previously published work measured H2A phosphorylation in pre-replicative rings exposed to ionising radiation (25), whereas we measured it in actively replicating trophozoites, wherein the background signal may be higher.) Therefore, we chose the replication-reduction assay as a more robust readout for checkpoint activity in subsequent experiments.

**Fig 1:**
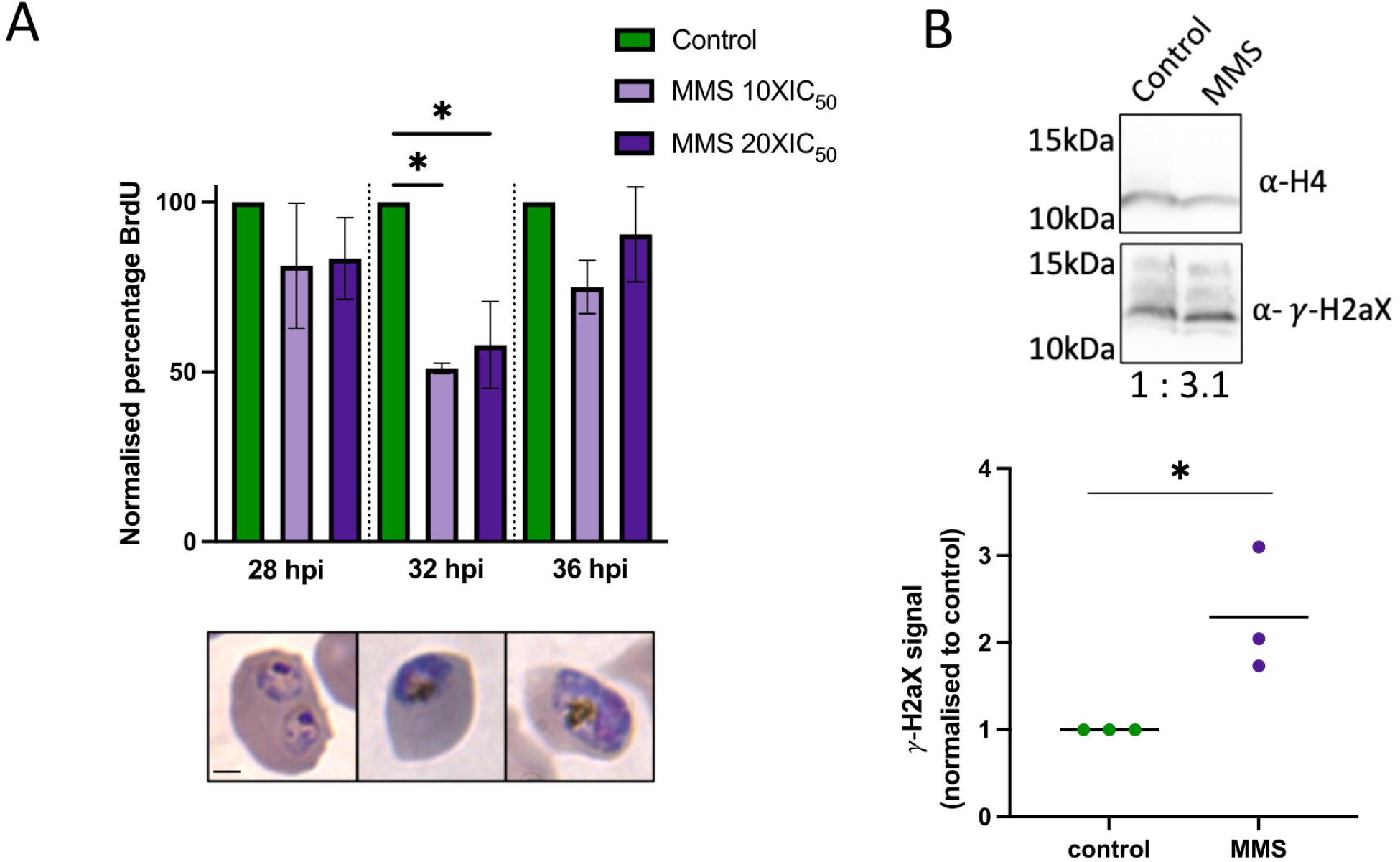
Evidence for DNA-damage-responsive checkpoint activity in *P. falciparum*. A. BrdU incorporation into populations of trophozoites at three stages, after treatment with MMS, as measured by ELISA. Parasites were treated with MMS for 1 h and BrdU was added for the latter 30 mins. Means of two biological replicates, conducted in technical triplicate, are shown; error bars show range. MMS treatments: 10x IC_50_, 960 µM; 20x IC_50_, 1.92 mM. For each parasite stage, statistical testing was via ordinary one-way ANOVA and posthoc Tukey’s multiple comparisons test. ^*^, P-value <0.05; comparisons not shown, ns. Pictures show examples of trophozoites at each timepoint. Scale bar, 2µm. B. Representative western blot of phosphorylated histone in trophozoite parasite lysate treated with MMS (2 h, 10x IC_50_). Phosphorylated histone signal was quantified versus control histone H4: relative quantification is shown below the blot. Graph shows quantifications from three biological replicate blots, relative to either H4 or Hsp70 as a control protein. Unpaired two-tailed t-test: P-value = 0.0354.

### Absence of clear checkpoint mediators in *P. falciparum*

When the intra-S-phase checkpoint is triggered in other eukaryotes, both histone phosphorylation and arrested DNA replication are transduced through PIKKs (24, 26). Since Figure 1 shows that these phenotypes also appear in *P. falciparum*, a transducing kinase should be present. However, orthologues of ATM/ATR are not found in the *P. falciparum* genome (6). To test for the presence or loss of such orthologues more thoroughly, we performed an extensive phylogenetic analysis of PIKK homologues throughout apicomplexans and other major eukaryotic lineages (Fig 2). In this analysis, the lipid kinases PI3K and PI4K formed an outgroup relative to all ATM/ATR-like proteins. In eukaryotes, PI3/4Ks are generally the closest relatives of PIKKs (31) and we did not seek to re-establish this (e.g. by surveying apicomplexan genomes for any other kinases resembling PIKKs), we simply sought to establish patterns of PIKK presence/loss in apicomplexans.

**Fig 2:**
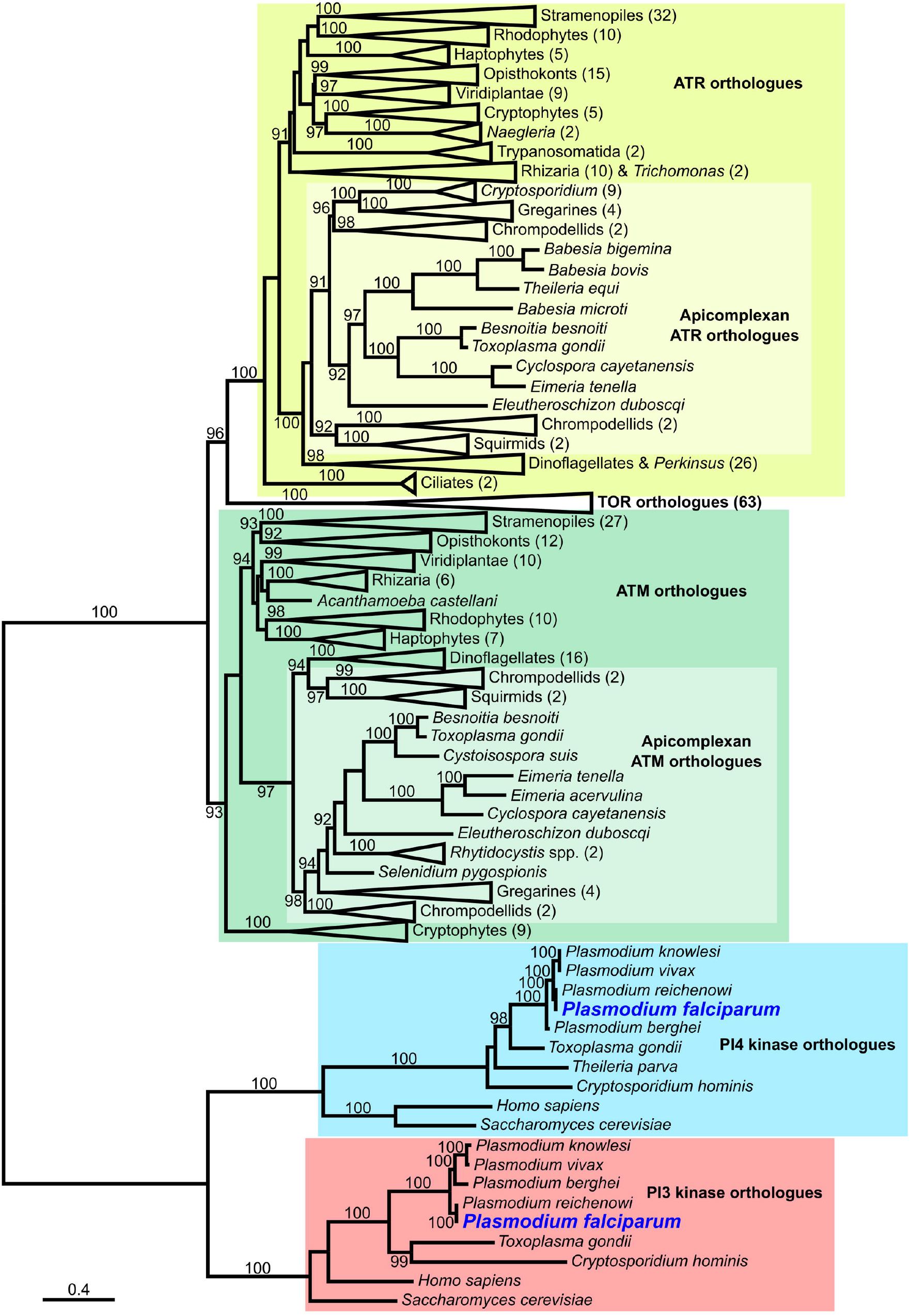
*Plasmodium* encodes no homologues of PIKK checkpoint kinases. C. A phylogeny inferred for amino acid protein sequences of ATR, ATM, and TOR homologues sampled from broad eukaryote groups. Apicomplexa homologues of PI3K and PI4K (the most closely related proteins to ATR and ATM present in *Plasmodium*) and characterised homologues of PI3K and PI4K from *Saccharomyces cerevisiae* and *Homo sapiens* were included as an outgroup. Support values, displayed as percentages, were generated from 1000 ultrafast bootstrap replicates (60) and only support values ≥ 90, indicating strongly supported clades, are shown. The scale bar represents the number of amino acid substitutions per site. A fully-expanded version of this tree with no collapsed clades and displaying sequence accessions is provided in the source data.

ATR homologues were found in most apicomplexans, and some – like *T. gondii* – clearly retained homologues of both ATM and ATR. Other apicomplexans, including *Cryptosporidium, Babesia* and *Theileria*, had independently lost the ATM homologue, indicating a propensity for loss of PIKKs in Apicomplexa. *Plasmodium* spp., however, had lost both ATM and ATR, retaining no apparent PIKKs at all. The closest PIKK homologues in any *Plasmodium* genome were the lipid kinases PI3K and PI4K (Fig 2, Fig S3), which were also present throughout apicomplexans (32). (*Theileria*, unusually, was found to have lost its PI3K homologue, possibly due to a lifecycle conducted free in the cytoplasm of leukocytes, which may supply this activity.

Notably, the closest *Plasmodium* PIKK homologue, PI3K, is reportedly overexpressed in artemisinin resistant parasites (33), which can alter their cell cycle dynamics. Furthermore, a recent analysis of the steady state locations of mature schizont-stage *P. falciparum* proteins by spatial proteomics assigned PI3K as a nucleus-located protein, whereas PI4K was assigned to cytoplasmic transport vesicles (Fig S4A) (34). This is consistent with PI3K having access to nuclear substrates such as histone H2A for phosphorylation. Accordingly we hypothesised that PI3K might moonlight as a checkpoint kinase in *Plasmodium*.

*Pf*PI3K is the only phosphoinositide 3-kinase of the six phosphoinositide kinases in *Plasmodium* (32): it is a well-validated, essential lipid kinase (35), with roles in nutrient uptake and vesicle trafficking (36, 37). However, its moonlighting as a protein kinase would not be unprecedented because other PI3Ks can to act on proteins as well as lipids. For example, humans encode a large family of PI3Ks, and their type-1 PI3Ks can phosphorylate protein substrates like cytokine receptors, as well as auto-phosphorylating themselves (38, 39). The sole *Plasmodium* enzyme is a type-3 PI3K, and thus far these are proven only to phosphorylate lipids (35).

We sought orthogonal validation of the spatial proteomics via immunofluorescence after raising peptide antibodies to *Pf*PI3K (Fig 3A, Fig S4B, C). The protein appeared in puncta throughout the cytoplasm, consistent with previous work (37) (Fig 3B). This highly dispersed location could obscure any minor nuclear signal, and we did not detect substantial re-location to nuclei upon DNA damage (Fig 3C). We then used expansion microscopy to better resolve the location of *Pf*PI3K, observing that puncta grew more abundant as cells matured, and were densest in an area free of nuclei that is probably the food vacuole – a known hub for vesicle trafficking (Fig 3D). Some *Pf*PI3K was also detected inside nuclei in all cells imaged (Fig 3E, Fig S5). Of note, we recently showed that the ATM and ATR homologues in *Toxoplasma* are likewise dispersed throughout the cell and do not substantially relocate to the nucleus after DNA damage, but they are nevertheless able to mediate checkpoint phenotypes such as histone phosphorylation (40).

**Fig 3:**
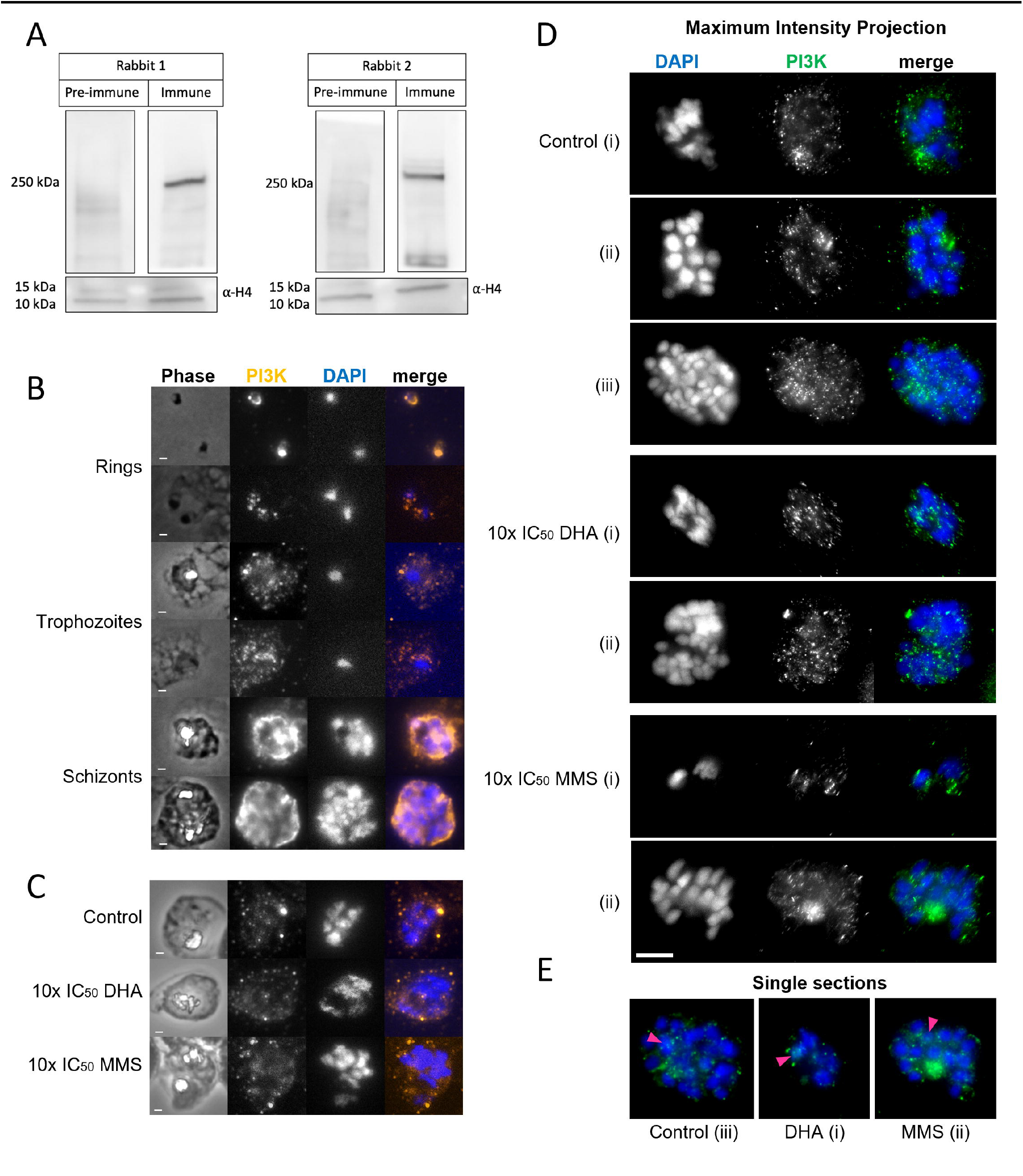
*Pf*PI3K is dispersed throughout the cell and does not visibly relocate to the nucleus upon DNA damage. A. Western blots showing detection of *Pf*PI3K by two antisera raised to *Pf*PI3K peptides. Replicate lanes from the same blot of protein lysate from late-stage *P. falciparum* were exposed to either pre-immune or immune serum. Histone H4 was used as a loading control. B. Immunofluorescence assay (IFA) showing location of *Pf*PI3K (detected by antiserum 1) in *P. falciparum*. 2 representative cells are shown per stage. Scale bar 1 μm. C. IFA showing *Pf*PI3K in *P. falciparum* after DNA damage: 1 h treatment with 5x or 10x IC_50_ of DHA or MMS on late-stage parasites. One representative cell per condition is shown. Scale bar 1μm. D. IFA showing *Pf*PI3K after expansion: maximum projections of 2-3 representative cells per condition, as in (C). DAPI, blue; *Pf*PI3K, green; scale bar 10μm. E. Single sections from the IFA in (D): nuclear *Pf*PI3K signals are highlighted with arrows. Full Z stacks are shown in supplementary data.

We attempted to use the same antibodies to immunoprecipitate native *Pf*PI3K with any potential protein substrates, but only a single *Pf*PI3K peptide was detected across several attempts. It is probably highly unstable *in vitro*, or lacking suitable tryptic digest sites, because it was likewise completely undetected, despite its very large size, in two dedicated proteomic studies of the nuclear versus cytoplasmic proteomes in *P. falciparum* (41, 42); and it was only sparsely detected in spatial proteomics (34). Further validation of an intra-nuclear role for *Pf*PI3K would probably require methods focussed less on the kinase itself and more on its targets – such as BioID for proximity-labelling of protein partners (43).

### The mediator of DNA-damage checkpoint responses in Plasmodium is sensitive to PIKK inhibitors

Assuming that a divergent PIKK-like kinase does exist in *P. falciparum*, we attempted to inhibit it with PIKK inhibitors. We chose two compounds with validated specificity for human ATM and ATR respectively (but not for human PI3Ks), KU-55933 and VE-821 (44, 45). If an ATM-like or ATR-like activity existed in *P. falciparum* – either in *Pf*PI3K or elsewhere – these compounds might specifically inhibit it.

We first determined their 48h IC_50_ values on *P. falciparum* (3D7 strain): KU-55933 had an IC_50_ in the micromolar range and VE-821, in the nanomolar range (Fig 4A, Fig S1B). Both inhibitors clearly synergised with MMS in a 48h isobologram analysis, particularly strongly in the case of VE-821, suggesting activity in the same pathway (Fig 4B). Control compounds that kill parasites but are not known to cause DNA damage – the antibiotics blasticidin and geneticin – did not synergise with MMS (Fig S6). Furthermore, when parasites were exposed to high-dose MMS (10x IC_50_, as in Figure 1) simultaneously with KU-55933 or VE-821, the inhibitors ablated the acute reduction in DNA replication caused by MMS, whereas co-exposure to the control antibiotic geneticin did not (Fig 4C). In this short 1h assay, we used the PIKK inhibitors at only their 48h IC_50_ levels, to minimise any negative effects from the inhibitors alone. Indeed, within the timeframe of the assay the inhibitors alone did not significantly affect DNA replication (Fig 4C). Overall, these data strongly supported the existence of a cryptic PIKK activity, responsible for a genuine intra-S-phase checkpoint via *in trans* kinase activity.

**Fig 4:**
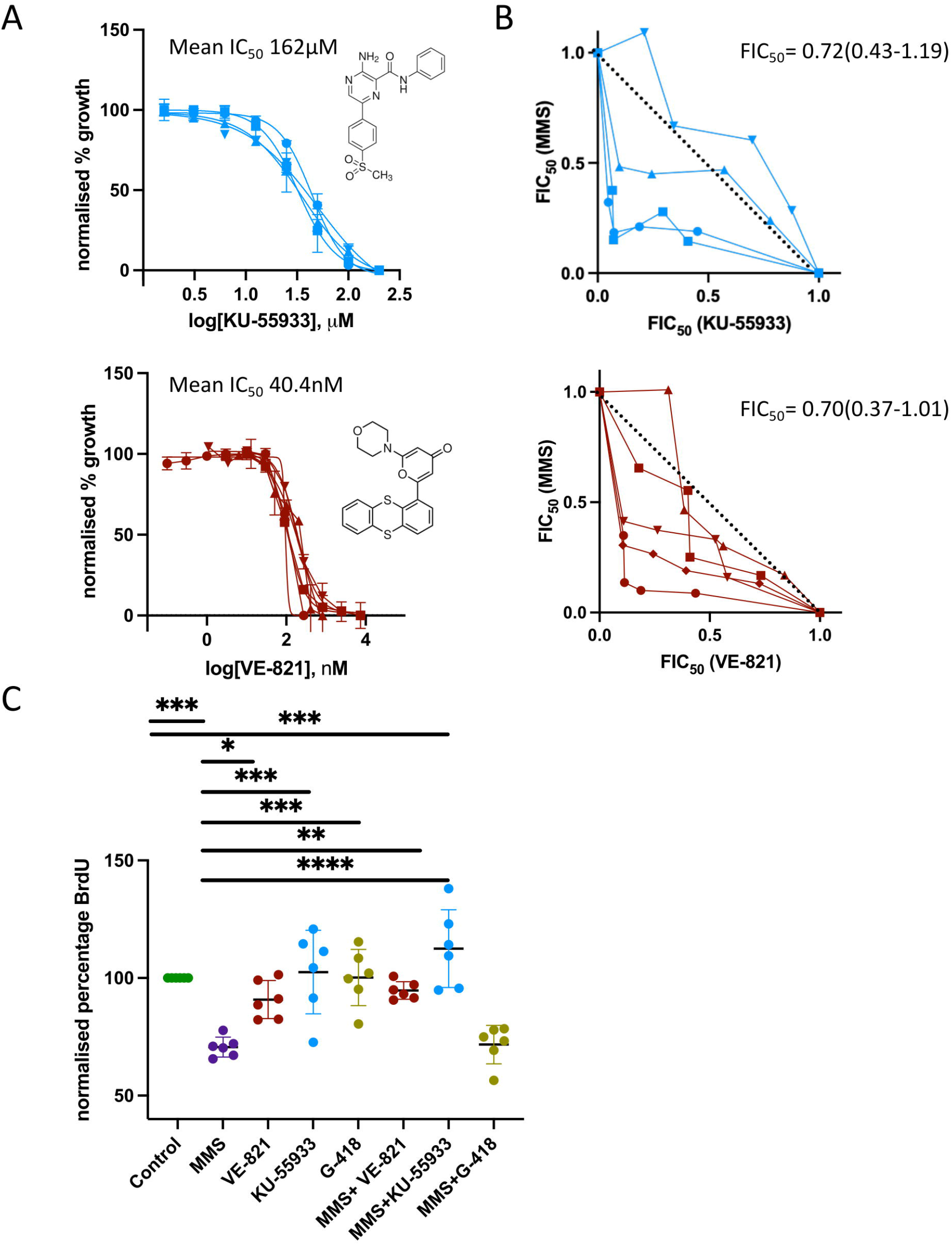
Inhibitors of human PIKKs synergise with the DNA-damaging agent MMS in *P. falciparum*. A. Inhibition of parasite growth by VE-821 and KU-55933 (molecular structures of each inhibitor are shown). Growth curves from five independent malaria SYBR green I-based fluorescence (MSF) assays: percentage parasite growth is plotted as a function of drug concentration, error bars are standard errors of the means. Mean IC_50_s from all experiments are shown (individual IC_50_ values from replicate experiments are in Fig S1B). B. Isobolograms plotted as fractional inhibitory concentrations (FIC) of combinations of MMS and VE-821 or KU-55933, as determined by MSF assay. FIC calculations were completed for each combination by dividing the measured “apparent” IC_50_ values for individual drugs in the different combinations by IC_50_ values obtained when the drugs were used alone. Each curve represents a different biological replicate experiment. The mean FIC (∑FIC(FIC_A_+FIC_B_)) is shown, with range. The type of relationship was defined as moderately synergistic based on well-established criteria: ∑FIC <0.5, substantial synergism; ∑FIC <1, moderate synergism; ∑FIC ≥1 and <2, additive interaction; ∑FIC ≥2 and <4, slight antagonism and ∑FIC >4, marked antagonism (61). C. BrdU incorporation into 30-32 hpi trophozoites after treatment with MMS, VE-821, KU-55933, geneticin and combinations of these, measured by ELISA. Parasites were treated with 10x IC_50_ of drug for 1 h and BrdU was added for the latter 30 mins. Six biological replicates are plotted, each completed in technical triplicate. Mean and standard deviation error bars are plotted, values normalised to control (set to 100%) in each replicate. Concentrations of drugs used: MMS 960 *µ*M, VE-821 1.62 *µ*M, KU-55933 400 µM, geneticin 1500 µg/ml. Ordinary one-way ANOVA and posthoc Tukey’s multiple comparisons test of control vs all and MMS vs all was completed; P-values: ^*^, <0.05; ^**^, <0.01; ^***^, <0.001; ^****^, <0.0001; comparisons not shown, ns.

### The antimalarial drug dihydroartemisinin induces DNA replication arrest in *P. falciparum*

MMS is a useful, well-characterised DNA-damaging agent, but it is not an antimalarial drug. We were particularly interested in the potential DNA-damaging activity of antimalarial drugs, because if parasites do make checkpoint responses to such drugs, then their antimalarial activity might be potentiated by inhibiting checkpoint kinases. Artemisinin is such a drug: it has non-specific alkylating activity that may affect both proteins and DNA (46, 47). There are similarities between the parasite’s response to dihydroartemisinin (DHA, the active metabolite of artemisinin) and MMS in terms of DNA breakage (46, 48), delay in the parasite’s transcriptional cascade, and upregulation of DNA repair genes (18). Indeed, we found that DHA induced H2A phosphorylation, at all stages of the cycle but particularly in pre-replicative ring stages (∼3-fold upregulation after 4h exposure to IC_50_ DHA, (Fig 5A)). This confirmed that DHA does indeed cause damage that activates a putative histone kinase, and furthermore that it can cause this damage prior to S-phase.

**Fig 5:**
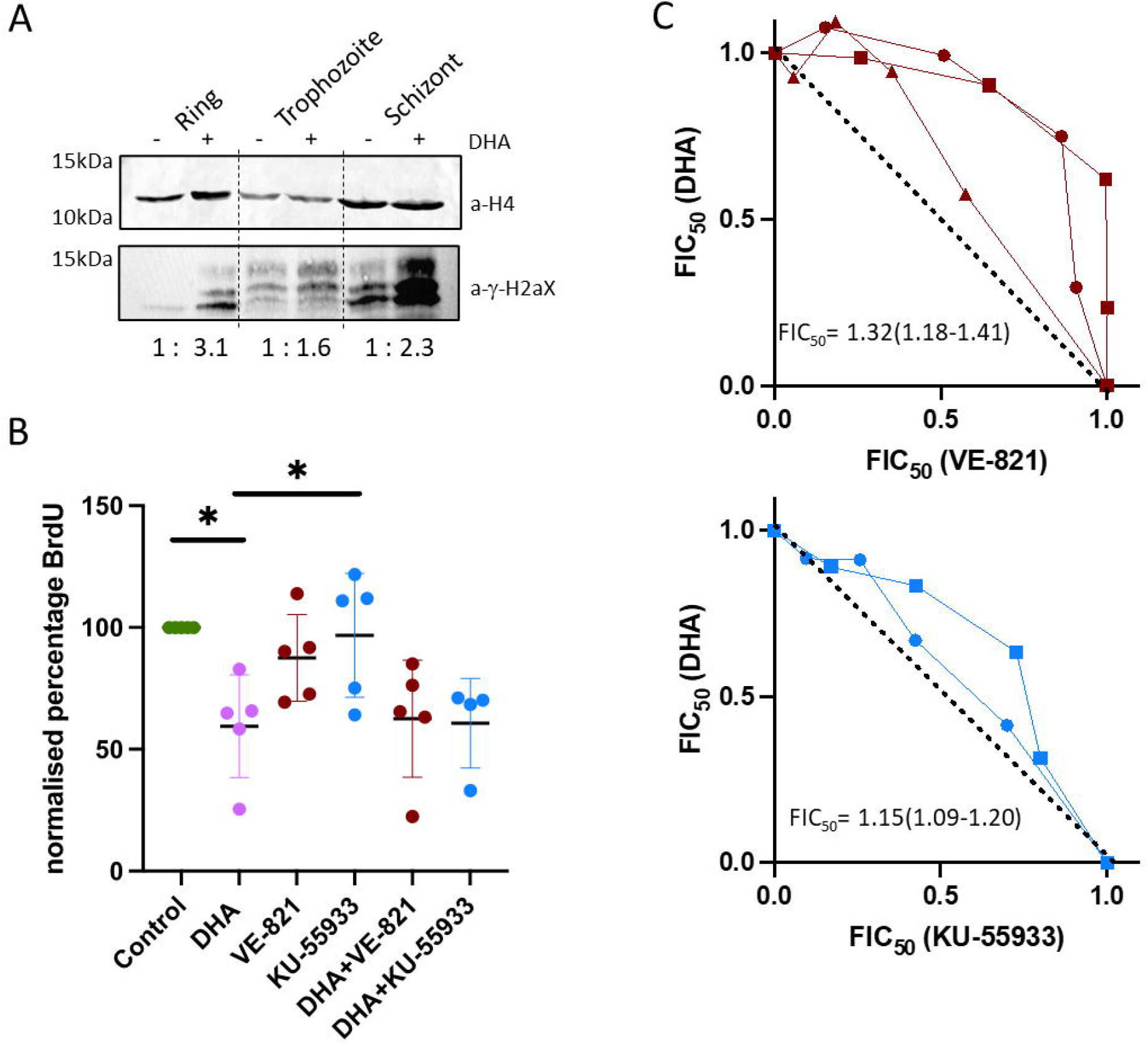
The antimalarial drug dihydroartemisinin induces a DNA replication arrest in *P. falciparum*. A. Western blot of phosphorylated histones in ring, trophozoite and schizont-stage parasites treated with DHA (4h, IC_50_ (3nM)). Phosphorylated histone signal was quantified versus control histone H4: relative quantification is shown below the blot. B. BrdU incorporation into 30-32 hpi trophozoites after treatment with DHA, VE-821, KU-55933 and combination treatments. Parasites were treated with 10X IC_50_ of drug for 1 h and BrdU was added for the latter 30 mins. Five biological replicates were plotted, each completed in technical triplicate. Mean and standard deviation error bars are plotted; values were normalised to control (set to 100%) in each replicate. Concentrations of drugs used: DHA 45 nM, VE-821 1.62 *µ*M, KU-55933 400 *µ*M. Statistical testing by ordinary one-way ANOVA and posthoc Tukey’s multiple comparisons test of control vs all and DHA vs all; P-values: ^*^, <0.05; comparisons not shown, ns. C. Isobolograms plotted as Figure 4B. Each curve is a biological replicate and the mean FIC (∑FIC(FIC_A_+FIC_B_)) is shown, with range. The relationship between DHA and either VE-821 or KU55933was defined as additive.

Using the replication-reduction assay, we saw that in replicating parasites, DHA caused an acute reduction in nascent DNA replication, similar to MMS (Fig 5B). However, unlike MMS, this was not ablated by the PIKK inhibitors KU-55933 and VE-821. Consistent with this difference, the inhibitors did not synergise with DHA in isobologram analysis (Fig 5C). This suggested that although both DHA and MMS may damage DNA, they differ in either the type of damage or the cellular response thereto in terms of inducing an intra-S-phase checkpoint.

### Ring-stage survival after artemisinin damage is affected by PIKK inhibitors

Cell-cycle checkpoint activity in *P. falciparum* may not be limited to an intra-S-phase checkpoint: most eukaryotic cells also operate a G1/S checkpoint, delaying S-phase entry if cells are damaged, starved or stressed. This may be analogous to the ‘TGA’ arrest or ‘dormancy’ that occurs in ring-stage parasites after artemisinin treatment (13). We therefore set out to establish whether ring-stage dormancy induced by DHA actually represents a classical, G1/S checkpoint, responsive to the DNA damage caused in ring stages by DHA (as shown in figure 5A), and mediated by a potential PIKK-like kinase.

We conducted the ring-stage survival assay (RSA) on parasites treated with DHA in the presence or absence of PIKK inhibitors. In this assay, pre-replicative early-ring-stage parasites are damaged with a pulse of DHA: most of them die, but some successfully arrest and thence survive. The more parasites are able to do this, the higher the subsequent recovery of viable parasites. We reasoned that if successful arrest is due to PIKK-mediated checkpoint activity, then PIKK inhibitors should reduce ring-stage survival. These experiments were conducted in the 3D7 strain and also in the Cambodian strain MRA-1252, derived from a naturally-arising DHA-resistant strain reverted back to a sensitive phenotype by specific correction of the resistance mutation in the *Kelch13* gene (49): its genetic background is otherwise that of a S.E. Asian DHA-resistant line.

Both 3D7 and MRA-1252 are highly sensitive to artemisinin, so we used the ‘extended recovery ring-stage survival assay’ (eRRSA), which detects recovered parasites very sensitively after 72h via quantitative PCR (50). Figure 6A shows that PIKK inhibitors, particularly KU-55933, did exacerbate the killing of ring-stage parasites by DHA. Differences were subtle and parallel assays reading the RSA with the less sensitive flow cytometric assessment of live/dead cells likewise showed marginal differences (Fig S7). However, in triplicate eRRSAs on two independent lines, 3D7 and MRA-1252, survival trended consistently downwards when DHA was combined with either inhibitor – so the effect, although not strong, was highly reproducible. As above, we minimised any negative effects of the inhibitors themselves by using them at a low level: only half of the 48h IC_50_. We confirmed that this exposure alone for 6h did not affect parasite survival after 72h (Fig 6B).

**Fig 6:**
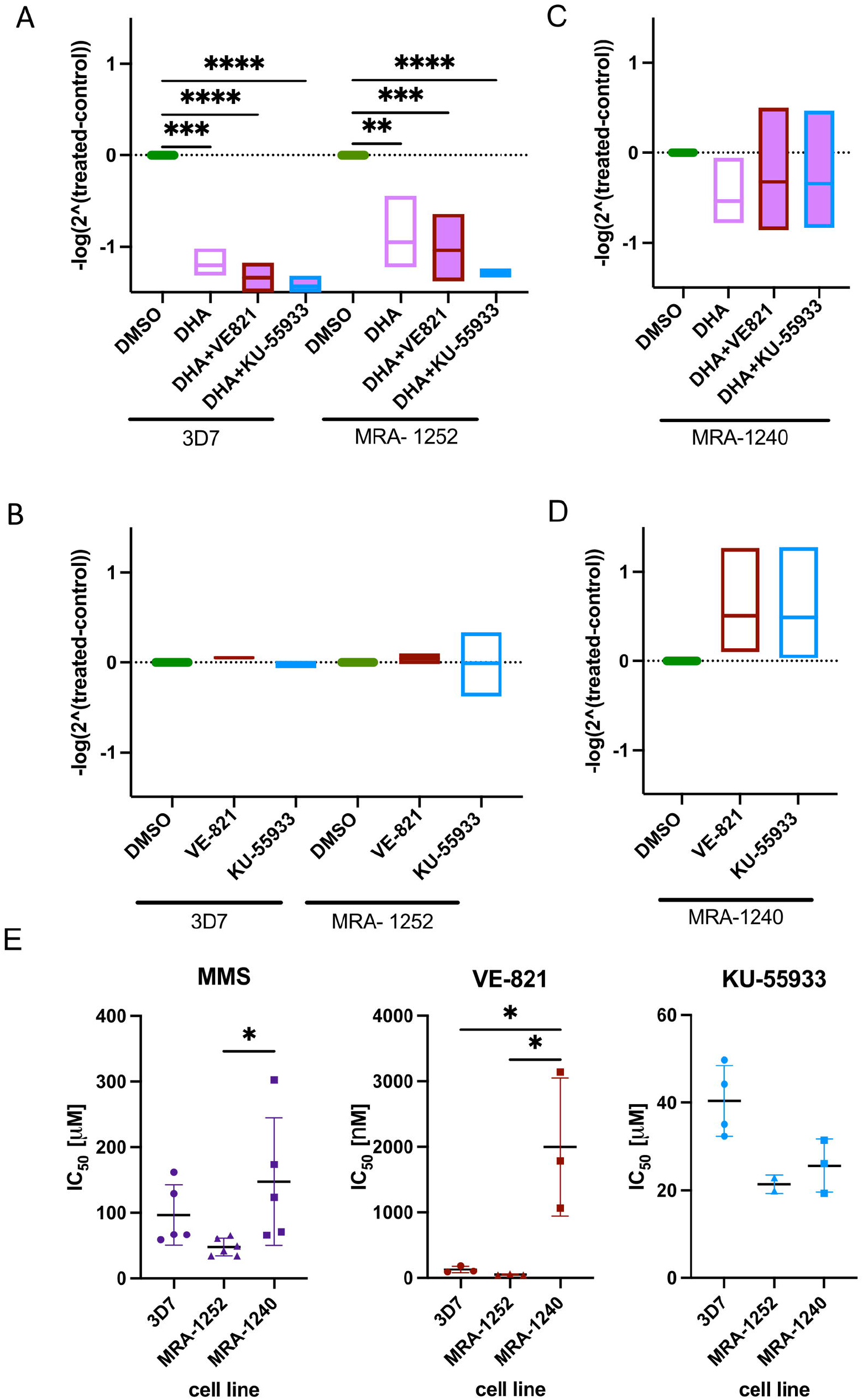
Ring-stage survival is affected by PIKK inhibitors. Data in A-D are from eRRSA assays on the strains 3D7, MRA-1252 and MRA-1240. Samples were treated with DMSO (the control condition), VE-821, KU-55933, DHA and combinations of these. Drug concentrations: VE-821, 81 nM (0.5X IC_50_); KU-55933, 20 μM (0.5X IC_50_); DMSO, 0.02%; DHA, 700 nM. Values plotted are the means of -log(foldchange = 2^(DHA-DMSO)) for each treatment. Midline represents the mean and boxes show min to max from three biological replicates, each conducted in technical triplicate. Statistical testing by ordinary one-way ANOVA and Dunnett’s multiple comparisons test, comparing all means to the mean value from DMSO- or DHA-treated samples. P-values: ^*^, <0.05; ^**^, <0.01; ^***^, <0.001; ^****^, <0.0001. Differences between DHA-treated samples +/-PIKK inhibitors did not reach statistical significance, but the p-values for DHA + VE-821 or DHA + KU-55933 versus control were < 0.0001, rather than 0.001 for DHA-alone versus control. A. Ring-stage DHA-sensitive parasite strains, 3D7 or MRA-1252, were treated with DMSO (control condition), DHA and inhibitors VE-821 or KU-55933 in combination with DHA for a 6 h pulse, then left to recover for 72 h. B. Control experiments without DHA: ring-stage 3D7 or MRA-1252 parasites were treated with DMSO or inhibitors VE-821 or KU-55933 for a 6 h pulse, then left to recover for 72 h. C. Ring-stages of the DHA-resistant parasite strain MRA-1240 were treated with DMSO (control condition), DHA and inhibitors VE-821 or KU-55933 in combination with DHA for a 6 h pulse, then left to recover for 72 h. D. Control experiment without DHA: ring-stage MRA-1240 parasites were treated with DMSO or inhibitors VE-821 or KU-55933 for a 6 h pulse, then left to recover for 72 h. E. Inhibition of growth over a full 48h growth cycle by MMS, VE-821 and KU55933 was measured in the strains 3D7, MRA-1252 and MRA-1240 via MSF assay. IC_50_s were calculated from at least three independent experiments and all values are plotted, with the midline showing the mean and error bars showing standard deviation. MSF growth curves are in Fig S1C.

These RSAs did not show conclusively that ‘TGA’ after artemisinin damage represents a true G1/S checkpoint but they did suggest that such a checkpoint, mediated by a cryptic PIKK, may contribute to ring-stage survival. Hence, the cryptic PIKK must be active throughout the cell cycle, not only in actively replicating trophozoites.

When the same assay was conducted on DHA-resistant parasites (the parent strain of MRA-1252, called MRA-1240) the result differed. As expected, DHA survival was much better in MRA-1240, and PIKK inhibitors had no negative effect on this survival (Fig 6C). There was even a trend towards better growth and DHA survival when MRA-1240 was exposed to PIKK inhibitors (Fig 6D). Hence, when a Kelch13 mutation was present, conferring DHA resistance, any contribution of checkpoint activity to parasite survival was apparently ablated.

### Artemisinin-resistant parasites are cross-resistant to MMS and VE-821, but not to KU-55933

We hypothesised that artemisinin-resistant parasites like MRA-1240 might not require ‘G1/S’ checkpoint activity to survive DHA damage because they are more efficient at DNA repair. Several mutations in DNA repair genes have been identified in resistant strains (47), although the pathway(s) leading to improved DNA repair are not fully characterised. We checked whether the DHA-resistant line MRA-1240 was cross-resistant to MMS, and its IC_50_ was indeed significantly higher than that of the matched sensitive line MRA-1252 (and also higher than that of 3D7) (Fig 6E, left panel; Fig S1C). Since the point mutation in Kelch13 is the only difference between the two Cambodian MRA lines, a genetic background of enhanced DNA repair cannot fully account for this.

Another explanation could be that MRA-1240 has hyper-active checkpoint activity due to Kelch13-associated over-expression of PI3K (33), and that this stimulates hyperactive DNA repair. This explanation *would* be consistent with the Kelch13 mutant and revertant having different sensitivities to MMS. If so, the Kelch13 mutant state would also be expected to correlate with reduced sensitivity to PIKK inhibitors. Indeed, MRA-1240 was much less sensitive to the VE-821 inhibitor (Fig 6E, middle panel; Fig S1C) – so much so that it was difficult to measure a consistent IC_50_ via MSF assay. Curiously, however, the same line was not less sensitive to KU-55933 (Fig 6E, right panel; Fig S1C). This was the first instance in which VE-821 and KU-55933 had shown a clear difference in their effects on *P. falciparum*. It may suggest that in the MRA-1240/1252 genetic background (perhaps unlike 3D7), more than one checkpoint protein is required, and that one of the inhibitors has a broader inhibitory spectrum than the other. For example, *Pf*PI3K may indeed moonlight as a checkpoint kinase, uniquely unregulated after Kelch13 mutation and uniquely inhibited by VE-821, whereas *Pf*PI4K and/or other ‘off-target’ kinases may also be needed when broader-spectrum damage induces more complex checkpoint responses, and KU-55933 may inhibit this broader kinase spectrum.

Validating these speculations would require challenging experiments, such as biochemical assays on recombinant PI3/4K proteins (which are ∼250kDa in size), or robust conditional manipulation of the two large, essential genes encoding *PfPI3K* and *PfPI4K* in parallel. We did make several attempts to tag the *PfPI3K* gene for conditional knockdown, but the enzyme’s active site is encoded at the extreme 3’ end and may become mis-folded after 3’ tagging, because all our attempts failed. Similar efforts by other groups to tag or manipulate this gene were likewise unsuccessful or gave no phenotype (51, 52), while a recently-published mutant that irreversibly deletes the 3’ end of the *PfPI3K* gene was lethal (53). The very large size of the gene (6.6kb) also precludes its overexpression or dominant-negative overexpression, as well as recombinant protein production for enzyme-activity assays. Overall, these large and essential genes proved near-intractable to molecular genetics. Thus, their putative PIKK-like activity was not conclusively evidenced or disproved, but the dual nuclear/cytoplasmic location of *Pf*PI3K does suggest that it could moonlight on nuclear substrates.

## Discussion

This work provides the first evidence that *Plasmodium* parasites during blood-stage schizogony can enforce an intra-S-phase checkpoint. The checkpoint is evidenced by an acute reduction in the rate of DNA replication and increased phosphorylation of histone H2A in a position analogous to the canonical DNA damage marker H2AX. *P. falciparum* was previously shown to slow its cell-cycle progression after DNA damage (18) but it was not clear if this occurred through cell-cycle checkpoint activity (partly because the *Plasmodium* system lacked validated DNA damage markers, signals or transducers thereof). Markers still remain limited – there are, for example, no cyclin markers for cell-cycle phases, and the H2A phosphorylation that was reported as a DNA damage marker in ring stages (25) is, in our hands, a consistent but low-dynamic-range marker in trophozoite stages, probably because background levels of H2A-P are higher in actively-replicating parasites. However, the ability to measure nascent DNA replication in *Plasmodium* (54) does now allow intra-S-phase checkpoint activity to be measured. Thus, we showed that DNA replication was acutely arrested by two distinct alkylating agents, MMS and the antimalarial drug DHA.

The transducer(s) of checkpoint activity in *Plasmodium* remain cryptic. *Plasmodium* spp. lack identifiable orthologues of PIKK kinases, in contrast to many apicomplexans including *T. gondii* (40). Nevertheless, such activity is apparently present, because the reduction in replication caused by MMS was ablated by well-validated inhibitors of human PIKKs. Hence it must be induced *in trans* by a kinase-mediated signalling cascade (also consistent with H2A phosphorylation), rather than occurring simply because alkylated DNA bases physically block the replisome. Interestingly, DHA differed from MMS in this respect: it also induced H2A phosphorylation, and it also affected DNA replication, but PIKK inhibitors did not significantly ameliorate this. Possibly the acute effect of DHA – which is a much larger molecule than MMS – *is* primarily to block replisomes via bulky DNA adducts (a documented effect of alkylating agents in human cells (23)). If such adducts cannot be quickly bypassed whether a checkpoint kinase is active or not, then PIKK inhibitors might not ameliorate the acute reduction in DNA replication.

Other, more complex, explanations are also possible for the difference in response to MMS versus DHA. For example, slow DNA replication can, in general, stimulate compensatory activation of extra replication origins (55), but when damage to DNA is sensed, checkpoint kinases act *in trans* to suppress such origin firing (23). If checkpoint kinases are inhibited, then extra origins may fire regardless, hence MMS-damaged cells may keep replicating and encountering more damage, so genome replication ultimately fails to complete. This would explain how MMS and PIKK inhibitors ultimately synergise in cell killing. Perhaps the broader-spectrum damage that DHA causes to proteins as well as DNA (46), and/or the oxidative stress that it generates (48) means that compensatory origin firing does not happen, whether a checkpoint is activated or not. Hence PIKK inhibitors would not synergise with DHA in cell killing. To evidence these rather complex hypotheses, precision measurement of DNA replication dynamics at single-molecule resolution after different types of damage will be needed. This has been achieved in human cells (23), and is now in development for *Plasmodium*.

Turning to the possibility of a pre-replicative G1/S checkpoint, the evidence for this being mediated by a PIKK-like kinase is tantalising albeit not conclusive. In the RSA, which measures DHA killing of pre-replicative rings, PIKK inhibitors exacerbated parasite killing. The effect was modest, so if a classical (kinase-enforced) G1/S checkpoint does exist, it is probably not the sole defining factor in survival of artemisinin-induced dormancy. Nevertheless, the trend was consistent in triplicate assays across two genetically-distinct DHA-sensitive strains, 3D7 and MRA-1252. Checkpoint kinase activity could therefore help to protect ring-stage cells, perhaps by detecting DNA damage, stimulating repair, and preventing S-phase entry until repair is complete. By contrast, a DHA-resistant strain showed no effect of PIKK inhibitors. This may be because it can already suppress DHA activation in early rings and thus protect itself from DNA damage; or alternatively, it may still sustain some damage, but be rescued by hyperactive DNA repair (47). Consistent with this latter hypothesis, it was recently shown that when *Pf*PI3K is inducibly knocked out in a DHA-resistant Kelch-13 mutant background, the parasites become less DHA resistant – perhaps because they can no longer resist or repair DHA-related DNA damage when the mediating kinase (*Pf*PI3K) is lost (53).

If there is indeed hyperactive DNA repair in the MRA-1240 line, it cannot be conferred per se by a DHA-resistant genetic background because the matched strain MRA-1252 was not cross-resistant to MMS, whereas MRA-1240 was. However, it could be because the Kelch13 mutation, uniquely present in MRA-1240, confers hyper-checkpoint activity, thus stimulating hyperactive DNA repair. In any case, these data raise the intriguing possibility that artemisinin drugs could synergise with checkpoint inhibitors in killing ring-stage parasites; the caveat being that this might only work well in sensitive, not resistant, strains.

It is an attractive possibility that *Pf*PI3K is the cryptic checkpoint kinase, particularly if Kelch13 mutant strains do contain more *Pf*PI3K (33). However, we were not yet able to prove or disprove this, owing to the extreme difficulty of manipulating the large, essential *PfPI3K* gene. Broad-spectrum inhibitors such as wortmannin were used in previous work (37), but these are slow-killing and unable to separate lipid-kinase from putative protein-kinase activity. We used PIKK inhibitors instead, hoping to isolate a putative protein kinase activity. The fact that our PIKK inhibitors did not synergise with DHA in MSF assays, whereas specific inhibitors of distinct human PI3Ks (i.e. lipid kinases) *have* been reported to do so (56), may suggest that this succeeded, and that VE-821 and KU-55933 do target a distinct activity in *Plasmodium*. Of course, they may affect irrelevant off-targets, i.e. multiple or unexpected kinases, or even non-kinases. As an example, a repurposed human-kinase inhibitor was recently deployed to target the PK6 kinase in *Plasmodium*, but it actually targeted haem detoxification as well (57). More likely, however, is that these inhibitors genuinely inhibit a *Plasmodium* checkpoint kinase – which may reside in *Pf*PI3K, *Pf*PI4K, or another radically divergent protein. *Plasmodium* contains many kinases whose roles are yet-unknown (58), and the *Plasmodium* cell cycle is driven by markedly divergent kinases such as CRK4 (59), so it is entirely possible that *Plasmodium* checkpoints are enforced by novel kinase(s) as well.

In conclusion, we present new evidence for measurable cell-cycle checkpoint activity in *Plasmodium*. We also raise the considerable challenge of unambiguously identifying its mediators in this divergent eukaryotic parasite.

## Methods

Full Methods as follows are provided as Supplementary material:

- Parasite culture & parasite strains
- Database sampling and phylogenetic analysis
- BrdU Enzyme-linked immunosorbent assay (ELISA)
- Immunofluorescence
- Immunofluorescence with expansion microscopy
- Malaria SYBR Green-1 Fluorescence (MSF) assay
- Isobolograms
- Ring-stage Survival Assay (RSA)
- RSA analysis: qPCR based & flow cytometry-based
- Antibody production
- Western blotting

## Acknowledgements

We thank Dr Holly Matthews for early work on this project, Dr Ibtissam Jabre for help with H2A-P blots, Prof Kai Wengelnik for sharing a PI3K inducible-knockout line, and Profs Marcus Lee and Paul Roepe for helpful early discussions.

This work was funded by the European Research Council (ERC) under the European Union’s Horizon 2020 research and innovation programme (ERC-2016-COG 725126 to CJM); a PhD studentship from the University of Cambridge Department of Pathology to MKJ; a Biotechnology and Biological Sciences Research Council (BBSRC) grant APP54272 to RFW; and a Gordon and Betty Moore Foundation Investigator Award to RFW (https://doi.org/10.37807/GBMF9194). The funders had no role in study design, data collection, interpretation or the decision to submit the work for publication.

## Supporting information

**Fig S1: Growth curves determining IC**_**50**_ **values for MMS, VE-821 and KU-55933 in the parasite strains 3D7, MRA-1252 and MRC-1240**.

Each panel shows growth curves from 3-5 independent malaria SYBR green I-based fluorescence (MSF) assays: percentage parasite growth is plotted as a function of drug concentration; error bars are standard errors of the means.

A. Inhibition of growth of 3D7 strain by MMS.

B. IC_50_ values from the growth curves shown in main figure 3A.

C. Inhibition of strains MRA-1252 and MRA-1240 by the drugs VE-821, KU55933 and MMS.

**Fig S2: 3D7 parasites survive a short pulse of MMS or DHA at the trophozoite stage, but their maturation is delayed and progeny number per schizont, reduced**.

A. Images were taken 12h after trophozoites at 30-32hpi had been exposed to a 1 h pulse of 10X IC_50_ MMS or DHA (2 independent experiments, ‘1’ and ‘2’). Images show that at this timepoint (i.e. 42-44hpi), cultures were dominated by a mixture of immature and mature schizonts, without clear evidence of parasite death in the damaged cultures. Parasites were then treated with 75nM ML10 for 10 h, to arrest mature schizonts before egress, and re-imaged. Mature schizonts then dominated in all cultures.

B. Parasite stages imaged in (A) were categorised as either: early rings, small rings, large rings, early trophozoites, mid trophozoites, late trophozoites, immature schizonts or mature schizonts (2 independent experiments, n ≥ 100 from each experiment). Differences between the proportions of stages in control vs DHA-treated vs MMS-treated cultures were tested via two-way ANOVA and posthoc Tukey’s multiple comparisons test. P-values are shown in a table below the graph: ^*^, <0.05; ^**^, <0.01; ^***^, <0.001. Stars of matched colours across rows denote a significant difference between those conditions (e.g. at 44hpi, there were still more immature schizonts after MMS treatment than in the control (pink stars); 10h later, this difference disappeared, with more mature and fewer immature schizonts (denoted by pairs of yellow and blue stars). The only other difference was in the small numbers of rings that escaped and reinvaded after prolonged arrest.

C. Merozoites per schizont were counted in each of the cultures shown in (A) (n = 26-30 per culture). Numbers of merozoites substantially increased in all cultures after the long ML10-arrest period, which allowed mature schizonts to accumulate even after the damage-induced delay seen in (B). A reduction in merozoite number was, however, still seen after MMS damage, although this was not significant after DHA damage.

**Fig S3: Protein sequence alignment of P. falciparum PI3K, PI4K and PIKKs from S. cerevisiae and human**.

MUSCLE (3.8) alignment of catalytic domains of *P. falciparum* PI3K and PI4K against human ATM, ATR, PI3K, and yeast Mec1 and Tel1. Colour key is ClustalW standard. Highlighted in the yellow boxes are the catalytic loops in ATR and ATM. Percentage identity matrix is shown below the alignment.

**Fig S4: Predicted cellular locations of PfPI3K and PfPI4K, and anti-PfPI3K antibody production**

**A**. Data plotted from (34), highlighting the clusters of *P. falciparum* proteins classified via spatial proteomics as nuclear (yellow) and vesicular (brown). *Pf*PI3K (encoded by gene PF3D7_0515300, red arrow)) is classified as nuclear, but falls at the boundary of the two clusters, whereas *Pf*PI4K (encoded by gene PF3D7_0509800, green arrow) is classified as vesicular and falls in the centre of the vesicular cluster.

B. Model made in i-TASSER (62), threading 1500 C-terminal residues of the 2133-amino acid *Pf*PI3K against human ATR. Protein coloured blue to red, N-terminal to C-terminal. Peptides used as antigens are highlighted in white and circled: amino acids 1359-1374 and 2011-2025 of full length *Pf*PI3K.

C. Controls for specificity of immune versus pre-immune antisera against *Pf*PI3K. Representative images of single cells are shown. Scale bar 1 µm. Primary antibodies used were rabbit antisera, pre-immune or immune, diluted 1:200 in 3% BSA.

**Fig S5: Immunofluorescence after expansion, showing the location of PfPI3K**

Examples of full Z-stacks taken during imaging of expanded cells: these stacks generated the maximum projections shown in Figure 3D for control cell (iii), DHA-treated cell (i) and MMS-treated cell (ii). They include the single sections extracted for Figure 3E. DAPI, blue; *Pf*PI3K, green; scale bar 10μm.

**Fig S6: Isobolograms showing that two antibiotic drugs do not synergise with MMS**

Isobolograms plotted as fractional inhibitory concentrations (FIC) of combinations of MMS and geneticin (G-418), or blasticidin (BSD), as determined by MSF assay. Data were processed as in Figure 4B. The type of relationship was defined as additive for blasticidin and borderline antagonistic for geneticin, based on well-established criteria (∑FIC <0.5, substantial synergism; ∑FIC <1, moderate synergism; ∑FIC ≥1 and <2, additive interaction; ∑FIC ≥2 and <4, slight antagonism and ∑FIC >4, marked antagonism).

**Fig S7: Ring stage survival assessed by flow cytometry**

A ring-stage survival assay was assessed by flow cytometry as well as eRRSA as used in Figure 5. Density plot quadrant graphs show one biological replicate of an RSA conducted on 3D7. Cell counts for each quadrant are shown. Percentage survival was calculated as the proportion of live versus dead cells (SYBR+ MITO+/SYBR+ MITO-) for each condition.

**Fig S8: Raw growth-curve data for all isobolograms plotted in figures 4B, 5C and S5**

A. Data for isobolgrams in figure 4B, VE-821 with MMS

B. Data for isobolgrams in figure 4B, KU-55933 with MMS

C. Data for isobolgrams in figure 5C, VE-821 and KU55933 with DHA

D. Data for isobolgrams in figure S5, Blasticidin and G-418 with MMS

**Source data files 1-6**

Files 1-6: Source data for phylogenetic tree shown in figure 2

