## Supplementary material for "Cell cycle checkpoint activity in the malaria parasite *Plasmodium falciparum*": Supp

### Methods

#### Parasite cultures

*Plasmodium falciparum* cultures were grown in 2% or 4% haematocrit human erythrocytes (Research Red Cells obtained from the NHS Blood & Transplant service), in RPMI 1640 (Sigma, R4130) supplemented with 2.3 g/L sodium bicarbonate, 50 mg/L hypoxanthine (Sigma, H9377), 25 µg/L gentamicin (Melford Laboratories, G38000-1), 2.5 g/L Albumax II and 5% human serum. Parasites were cultured under a 1% $O_2$/ 3% $CO_2$/ 96% $N_2$ gaseous atmosphere at 37 °C.

#### Parasite strains

Parasite strains used were: 3D7, 3D7 pTK, iPI3K-ko (also in the 3D7 background, gift from Prof Wengelnik) [1], MRA-1240 and MRA-1252. 3D7 pTK carries a thymidine kinase transgene to allow incorporation of bromodeoxyuridine (BrdU) for monitoring DNA replication [2]. The iPI3K-ko line carries a D-Cre-inducible deletion module for the C-terminal part of the *PI3K* gene [1]. MRA-1240 and MRA-1252 are matched artemisinin-resistant and sensitive Cambodian field lines, respectively [3], obtained through BEI Resources (NIAID, NIH). Parasites were synchronised as in [4].

#### Database sampling and phylogenetic analysis

The phylogeny (Fig 2) for ATM, ATR, TOR, PI3K and PI4K homologues was inferred by following a computational workflow (https://github.com/camwallerlab/Methods-for-phylogenetic-analysis-of-plastid-translocons) developed as part of a previous study [5]. In summary, this workflow involved sampling a custom protein sequence database that included data for major eukaryotic groups from UniProt [6] and the Marine Microbial Eukaryote Transcriptome Sequencing Project [7], as well as data for apicomplexans and close relatives from VEuPathDB [8] and two previous studies [9, 10]. This custom database was then sampled for ATM, ATR and TOR homologues using the program blastp (BLAST+ version 2.11.0 [11]), for which characterised homologues of these proteins from selected model organisms were used as search queries. The dataset of possible ATM, ATR and TOR homologues obtained was then clustered with CD-HIT [12] to remove highly similar sequences, thereby reducing overall redundancy of the dataset. PI3K and PI4K homologues from *Plasmodium* and other apicomplexans, as well as characterised homologues from *Saccharomyces cerevisiae* and *Homo sapiens* were incorporated into the dataset as a phylogenetic outgroup for the ATM, ATR and TOR proteins. Iterative rounds of alignment were then performed using mafft (MAFFT version 7.475 [13]), conserved site selection using trimal with "gappyout" mode (trimAl version 1.4 [14]), and tree inference with FastTreeMP (FastTree version 2.1.11 [15]) using the default settings. For each iterlation, sequences that aligned poorly or were so dissimilar from the rest of the sequences in the dataset that they were unlikely to be ATM, ATR or TOR homologues, but rather false positives of the described sampling strategy, were manually identified and removed from the dataset. The final curated dataset that these iterations obtained was then aligned using mafft-linsi (MAFFT version 7.475 [13]) and conserved sites were selected using trimal with "gappyout" mode (trimAl version 1.4 [14]). A phylogeny was then inferred from this dataset using the program iqtree2 (IQ-TREE version 2.1.2 [16]) with 1000 ultrafast bootstrap replicates (UFBoot2 [17]) and using the best-fitting model, LG+F+I+G4 ([18]) that was chosen according to the Bayesian Information Criterion by ModelFinder [19], all implemented within iqtree2. The protein sequences, alignment, and tree inference output files used to generate the phylogeny are provided as source data.

#### BrdU Enzyme-linked immunosorbent assay (ELISA)

ELISA was carried out as per [2]. In brief, tightly synchronised *P. falciparum* 3D7 pTK parasites were collected at 4% parasitaemia and 4% haematocrit at 28 hpi, 32 hpi and 36 hpi. They were labelled with BrdU (100 µM, Sigma) for 30 min and then collected for ELISA, or treated with MMS at 10 x $IC_{50}$ (960 µM) or 20 x $IC_{50}$ (1,920 µM) for 30 min prior to BrdU labelling. In further experiments, parasites at 30-32 hpi were exposed to either MMS alone (10x $IC_{50}$), MMS+VE-821 (1x $IC_{50}$), MMS+KU-55933 (1x $IC_{50}$), VE-821 alone, KU-55933 alone, DHA alone (10x $IC_{50}$), DHA+VE-821, DHA+KU-55933 or no drugs, then processed as above.

### Immunofluorescence

Air-dried thick blood smears were fixed with 4% paraformaldehyde in PBS for 10 min and permeabilised for 10 min with 0.05% triton in PBS. Slides were then blocked for 1 h in 1% BSA/PBS. The primary antibody (1:200 immune or preimmune antiserum) was added in blocking buffer for 1 h under a coverslip. After 3x 5 min washes in PBS, the secondary antibody was added to slides for 1 h. The slides were washed with PBS for 5 min, 120 µl of 2 µg/ml DAPI solution was added for 10 min and then the slides washed again with PBS for 5 min. Coverslips were mounted on slides in Prolong® Diamond (Invitrogen) and allowed to dry overnight at 4 °C. Images were acquired using a Nikon Eclipse Ti widefield microscope with a Nikon objective lens (Plan APO, 100x/1.45 oil), and a Hamamatsu C11440, ORCA Flash 4.0 camera. Images were processed using the NIS-Elements software and ImageJ (v.1.51w).

### Immunofluorescence with expansion microscopy

Expansion Microscopy was performed according to Liffner *et al.* [20].
12 mm round coverslips (Fisher, Cat# NC1129240) were treated with poly-d-lysine (1 h, 37°C), washed twice with MilliQ water, and placed in the wells of a 12-well plate. 1 mL of parasite culture was added to the well containing the coverslip for 15 min at 37°C. Culture supernatants were removed, and parasites were fixed with 1 mL of 4% v/v paraformaldehyde in PBS for 15 min at 37°C. Following fixation, coverslips were washed three times at 37°C with PBS before being treated with 1 mL of 1.4 % v/v formaldehyde/2% v/v acrylamide (FA/AA) in PBS. Samples were then incubated at 37°C overnight.

Monomer solution (19% w/w sodium acrylate [Sigma, Cat# 408220], 10% v/v acrylamide [Sigma, Cat# A4058, St. Louis, MO], 0.1% v/v N,N'-methylenebisacrylamide [Sigma, Cat# M1533] in PBS) was typically made the night before gelation and stored at −20°C overnight. Prior to gelation, FA/AA solution was removed from coverslips, which were washed once in PBS. For gelation, 5 µL of 10% v/v tetraethylenediamine (TEMED; Thermo Fisher, Cat# 17919) and 5 µL of 10% w/v ammonium persulfate (APS; Thermo Fisher, Cat# 17874) were added to 90 µL of monomer solution and briefly vortexed, then 35 µL was pipetted onto parafilm and coverslips were placed (cell side down) on top. Gels were incubated at 37°C for 30 min before transfer to wells of a 6-well plate containing denaturation buffer (200 mM sodium dodecyl sulfate (SDS), 200 mM NaCl, 50 mM Tris, pH 9). Gels were incubated in denaturation buffer with shaking for 15 min, before separated gels were transferred to 1.5 mL tubes containing denaturation buffer. The tubes were incubated at 95°C for 90 min. Following denaturation, gels were transferred to 10 cm Petri dishes containing 25 mL of MilliQ water for the first round of expansion and placed onto a shaker for 2x 15 min, changing water in between. Gels were subsequently shrunk with 2x 15 min washes in 25 mL of 1× PBS, before being transferred to 6-well plates for 30 min of blocking in 3% BSA-PBS at room temperature.

After blocking, gels were incubated with primary antibodies (1:200 immune antiserum), diluted in 3% BSA-PBS, overnight. Gels were washed three times in PBS for 10 min before incubation with secondary antibodies (1:1000) and DAPI diluted in 3% BSA-PBS for 3 h. Gels were again washed three times in PBS, before being transferred back to 10 cm Petri dishes for re-expansion with three 30 min MilliQ water incubations.

Gels were either imaged immediately following re-expansion or stored in MilliQ water until imaging, using a Nikon Eclipse Ti2 widefield microscope with a Nikon objective lens (Plan APO,

60x/1.40 oil), and a Hamamatsu C11440, ORCA Flash 4.0 camera. Z-stacks were taken with a depth range of 2-10μm, at interval range of 0.2-0.6μm (depending on the parasite stage). Images were processed using the NIS-Elements software and ImageJ (v.1.51w).

#### Malaria SYBR Green-1 Fluorescence (MSF) assay

MSF assays were conducted as in [21]. 100 μM chloroquine (Merck) was used as a control for parasite death. Data were analysed and graphs plotted in GraphPad Prism.

#### Isobolograms

Parasites were prepared as for MSF assays. Isobologram design followed the procedure in [22]. Serial dilutions were planned so that the $IC_{50}$s of each drug would be in the second two-fold dilution out of 6 total dilutions. $IC_{50}$s for each combination solution were calculated using sigmoidal best-fit curves assigned in Graphpad Prism. Fractional inhibitory concentrations (FIC values) were calculated in Excel [23] and isobolograms plotted in Graphpad Prism. Source data for the isobolograms are provided in figure S8.

#### Ring-stage Survival Assay (RSA)

Parasite cultures at 8-10% schizonts were synchronised using percoll and Compound 2 as in [4], then allowed to reinvade. Early ring-stage (0-3 hpi) parasites at 0.5% parasitaemia, 2% haematocrit were exposed for 6 h to 700 nM DHA or 0.02% DMSO (vehicle control) under normal conditions, before washing three times with incomplete media, resuspending in complete media and incubating under the same conditions for a further 66 h. In addition to the DHA 700 nM pulse, VE-821 and KU-55933 inhibitors were used at 0.5x $IC_{50}$ with DHA, or by themselves, throughout the 6 h pulse period. 72 h after initial drug exposure, 20 μl was frozen for RT-PCR as per eRRSA protocol [24].

#### qPCR-based RSA analysis

qPCR amplification was used to quantify live parasites in the eRRSA [24]. The Phusion Blood Direct PCR kit was used (ThermoFisher, cat # F547L), supplemented with 1x SYBR Green I (ThermoFisher) and 50 nM Low ROX (ThermoFisher), according to the manufacturer's instructions. qPCR amplification was measured using the fast mode of the ABI 7900HT, with a 20 s denaturation at 95 °C, followed by 35 cycles of 95 °C for 1 s, 62.3 °C for 30 s, and 65 °C for 15 s. Cycle threshold (Ct) values were calculated; then the fold change was calculated by determining the mean difference in Ct (ΔCt) for the three technical replicates between the untreated and DHA-treated samples by applying the following equation: Fold change =2^(ΔCt of treated – ΔCt of untreated). This was -log transformed and data were plotted in GraphPad Prism.

#### Flow cytometry-based RSA analysis

Flow cytometry wash buffer 'FC wash' (2% FBS (Gibco) in HBSS (calcium, magnesium, no phenol red, Gibco) was prepared. The remaining 180 μl of each RSA assay, not harvested for eRRSA, was centrifuged and packed cells were washed three times in 200 μL of FC wash, then resuspended in FC wash to make a 10% haematocrit suspension. 25 μL of suspension from each assay (conducted in technical triplicate) were mixed with 25 μL of 0.6X SYBR Green I (SYBR) with 1.5 nM MitoTracker Deep Red FM (MT, Invitrogen) in FC wash. Compensation controls were prepared with suspension from one DMSO-treated triplicate, labelled with either 0.6X SYBR, 1.5 nM MT, or FC wash alone.

Samples were incubated in the dark at 37°C for 30 min, then washed three times in 200 µL and resuspended in 180 µL of FC wash.

10,000 events of SYBR positive cells were captured at a flow rate of 25 µL/min on an Attune NxT Acoustic Focusing Cytometer (Invitrogen). SYBR and MT fluorescence were detected on the BL1 (488 nm excitation, 530/30 nm emission, 340 V) and RL1 (638 nm excitation, 670/14 nm emission, 510 V) channels respectively. Debris and doublets were excluded by gating on a forward scatter area (FSC-A) versus side scatter height (SSC-H) plot and forward scatter area (FSC-A) versus forward scatter height (FSC-H). SYBR/MT quadrant plots were generated using the Attune NxT software v4.2.0 (Invitrogen).

#### Antibody production

Peptides were designed from *P. falciparum* PI3K (PF3D7_0515300) with help from Eurogentec using parameters such as size, amino acid composition, hydrophobicity and secondary structure. Two peptides were chosen - one predicted to be in the catalytic domain and one upstream of this domain, both predicted to be surface-displayed. N-terminal cysteines were added to the peptide sequences to target the coupling site at the carrier protein.
Antibodies were raised by Eurogentec. Two rabbits were bled for pre-immune sera, inoculated with both peptides, then further immunised with 3 booster inoculations at 12 days, ~1 month and ~2months. Immune serum was collected after ~9 weeks.

#### Western blotting

Parasites were extracted from infected erythrocytes by mixing with 0.5 volumes of 0.2% saponin in PBS and incubating for 10 min on ice. Saponin lysates were centrifuged at 15,000 *xg* at 4 °C for 10 min, washed 3 x in ice-cold PBS, centrifuged at 15,000 *xg* for 5 min and resuspended in PBS before freezing at -20°C. Parasites obtained by saponin lysis were resuspended in 1x RIPA buffer (150 mM sodium chloride, 1.0% NP-40 or Triton X-100, 0.5% sodium deoxycholate, 0.1% SDS and 50 mM Tris pH 8) with protease inhibitors (ROCHE, Complete Mini) added directly before use, then freeze-thawed 3 times. Cell debris was removed by centrifugation at 4 °C, 13,000 *xg*, and supernatant was mixed with 1 x NuPage LDS Sample Buffer (ThermoFisher: NP0007), supplemented with 0.1% 2-mercaptoethanol, then boiled at 90 °C for 5 min. Protein lysates were resolved on 4-15% TRIS-glycine gels (BIO-RAD Mini -PROTEAN TGX) for 1–2 h at 100 V in 1X TGS buffer. Wet transfer was completed onto nitrocellulose membrane 0.2-µM-pore-size (Amersham Protran®, GE Healthcare) at 60 V for 45 min.

Membranes were blocked with 3% BSA TBST for 1 h, washed once in TBST and incubated overnight in primary antibody diluted in blocking buffer at 4 °C. Blots were washed 3x 5 min in TBST and incubated in a secondary antibody diluted 1:5000 in blocking buffer for 1 h. Blots were washed again (3x 5 min, TBST), developed using SuperSignal West Pico Plus Chemiluminescent Substrate and imaged on a gel imager (Azure, Q500). Some membranes were stripped prior to re-blocking (10 ml 20% SDS and 12.5 ml 0.5M Tris HCL, pH 6.8, added to 77.5 ml distilled water, with 0.8 ml β-mercaptoethanol added just before use). 5 ml stripping buffer (warmed to 50 °C) was added to a blot and incubated at 50 °C for 45 min with agitation. The stripping buffer was removed, and the membrane was washed under a running tap for 1-2 min, then washed 3x 5 min with TBST. The membrane was then ready for re-blocking.

Antibodies used were pre-immune or immune sera for *Pf*PI3K peptides, diluted 1:200, with secondary goat anti-rabbit HRP 1:5000 (Abcam, ab97080); or rabbit α-γ-H2aX (Cell signalling, 9718S) with goat α-Rabbit IgG (H+L) Cross-Adsorbed Secondary Antibody, Alexa Fluor 594 (Invitrogen, A-11012).

### Figure S1:

A

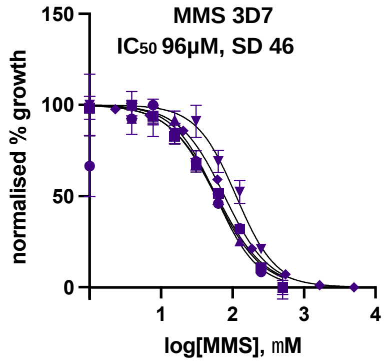


B

| | VE-821 [nM] | KU-55933 [$\mu$M] |
|---|---|---|
| $IC_{50}$ | 95.49 | 44.3 |
| | 106.7 | 32.31 |
| | 253.5 | 35.1 |
| | 190.4 | 49.8 |
| Mean $IC_{50}$ | 162 | 40.4 |

C

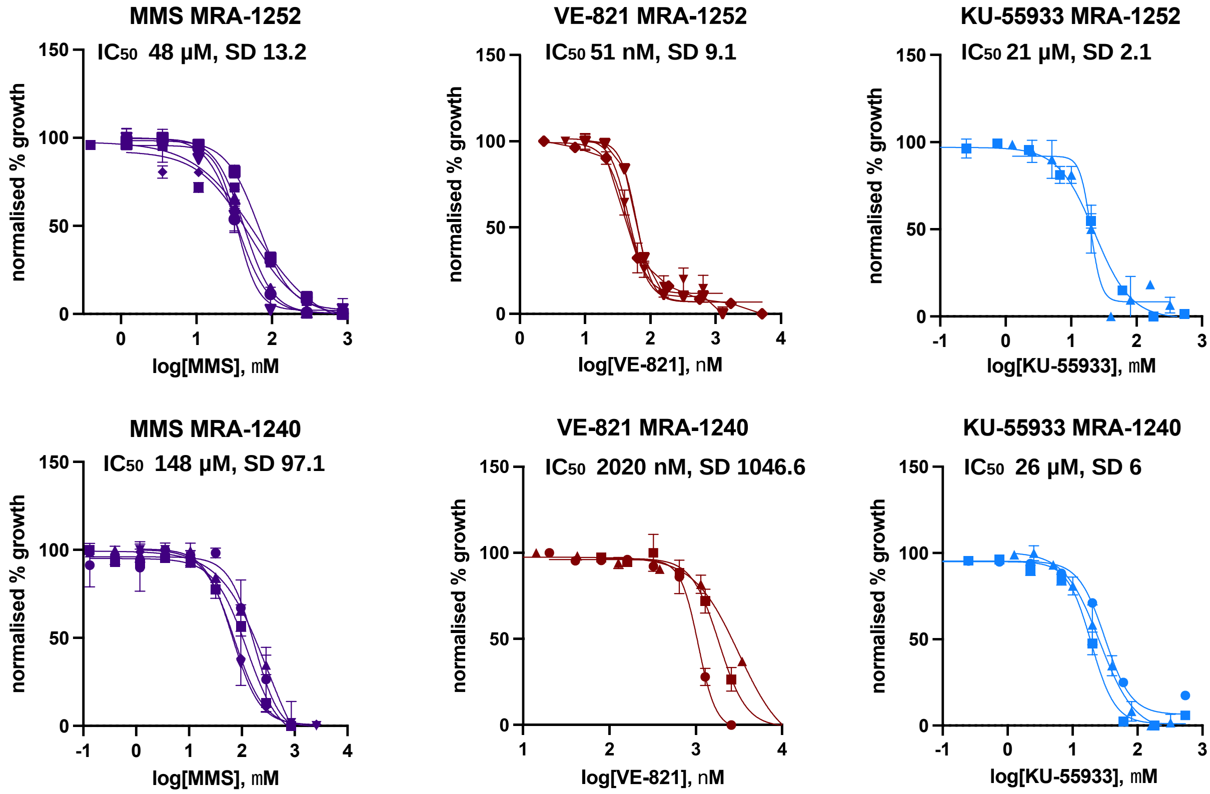

Figure S2:

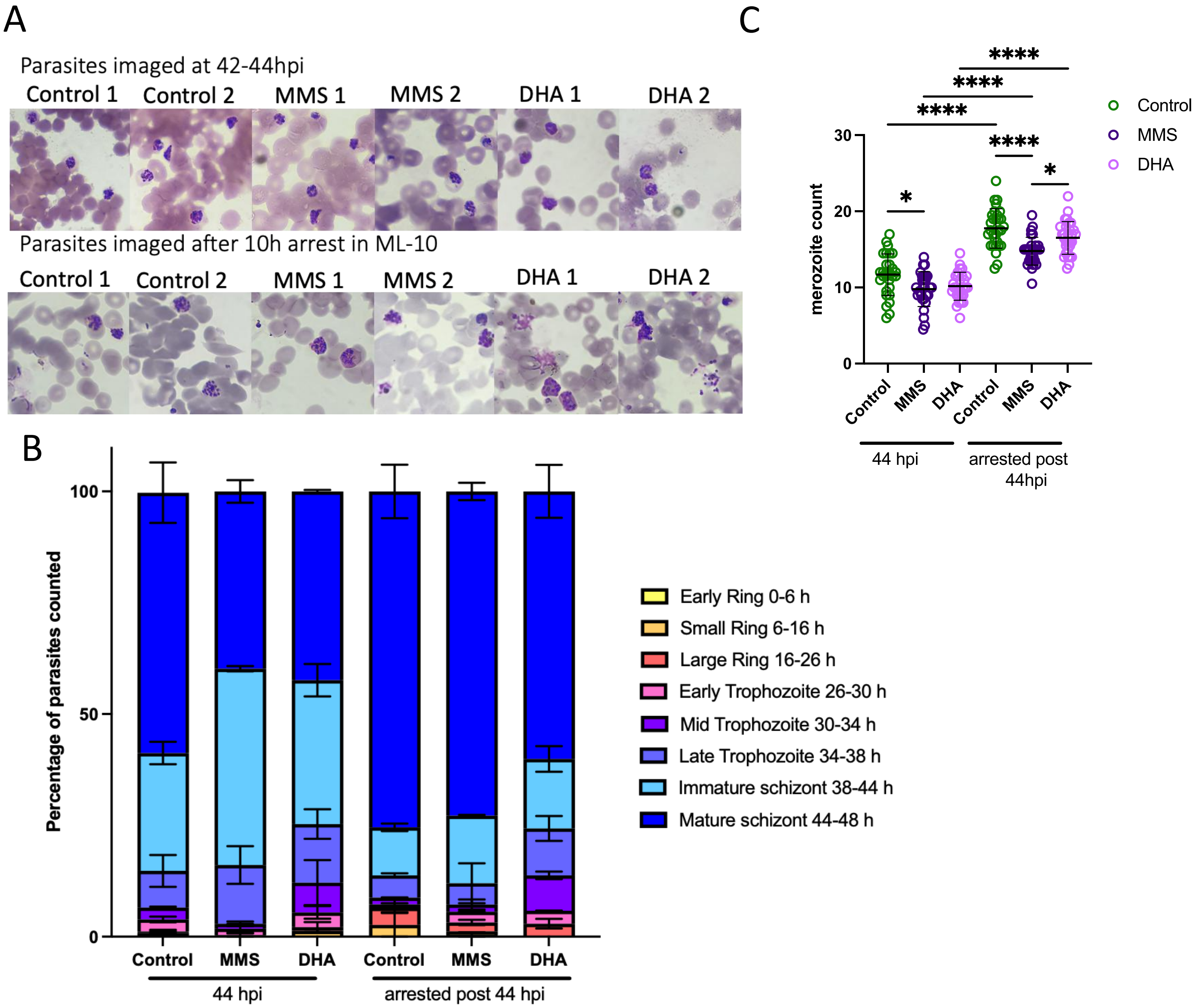


| 44 hpi | | | arrested post 44 hpi | | | |
|---|---|---|---|---|---|---|
| Control | MMS | DHA | Control | MMS | DHA | |
| – | – | – | – | – | – | Early Ring 0-6 h |
| – | * | – | – | * \| * | * | Small Ring 6-16 h |
| – | – | – | – | – | – | Large Ring 16-26 h |
| – | – | – | – | – | – | Early Trophozoite 26-30 h |
| – | – | – | – | – | – | Mid Trophozoite 30-34 h |
| – | – | – | – | – | – | Late trophozoite 34-38 h |
| *** | *** \| * | – | – | * | – | Immature schizont 38-44 h |
| – | * | – | – | * | – | Mature schizont 44-48 h |

### Figure S3:

LEGEND

A R N D C Q E G H I L K M F P S T W Y V B X Z

```
sp|Q13315|ATM_HUMAN                 FKAEFRLAGGVNLPKIIDCVGSDGKERRQLVKG-RDDLRQDAVMQQVFQMCNTLLQRNTE  2684
sp|P38110|TEL1_YEAST                NETVGITTTGLSLPKIVTFNISDGTTQKALMKGSNDDLRQDAIMEQVFQQVNKVLQNDKV  2943
sp|Q13535|ATR_HUMAN                 FDDMVEILASLQKPKKISLKGSDGKFYIMMCKP-KDDLRKDCRLMEFNSLINKCLRKDAE  2354
sp|P38111|MEC1_YEAST                FGSSYKVFSSLKKPKQLNIIGSDGNIYGIMCKK--EDVRQDNQYMQFATTMDFLLSKDIA  2106
PI4K_Plasmodium_falciparum_         ---------------------------VIIKG-GDDLRQELLASQLIKQFKIIFENA--  1334
tr|Q8I3V5|PI3K_Plasmodium_falciparum_ --------------------------KYIYKA-GDDLRQDHLVIQIIYVMDNIWKRY--  1907
sp|Q8NEB9|PI3KC3_HUMAN              IPETATLFKSALMPAQLFFKTEDGGKYPVIFKH-GDDLRQDQLILQIISLMDKLLRKE--  662
                                                               : *   :*:*::    :.    .    .

sp|Q13315|ATM_HUMAN                 TRKRKLTICTYKVVPLSQRSGVLEWCTGTVPIGEFLVNNEDGAHKRYRPNDFSAFQCQKK  2744
sp|P38110|TEL1_YEAST                LRNLDLGIRTYKVVPLGPKAGIIEFVANSTSLHQILSKLHTND-------KITFDQARKG  2546
sp|Q13535|ATR_HUMAN                 SRRRELHIRTYAVIPLNDECGIIEWVNNTAGLRPILTKLYKEKGVYMTGKEL-----RQC  2409
sp|P38111|MEC1_YEAST                SRKRSLGINIYSVLSLREDCGILEMVPNVVTLRSILSTKYESLKIKYSLKSL-----HDR  2161
PI4K_Plasmodium_falciparum_         --GLPLWLRPYEILVTGSNSGIIEYVNDTCSVDSLKRKF-GAD-------S---------  1374
tr|Q8I3V5|PI3K_Plasmodium_falciparum_ --GLDLKMTLYRVLALSTDDGFIEFVDYAESISSIKKNY-KGE-------I---------  1947
sp|Q8NEB9|PI3KC3_HUMAN              --NLDLKLTPYKVLATSTKHGFMQFIQSVPVAEV--LDT-EGS-------I---------  700
                                          * :  * ::      *.::

sp|Q13315|ATM_HUMAN                 MMEVQKKSFEEKYEVF-MDVCQNFQPVFRYFCMEKFL----DPAIWFEKRLAYTRSVATS  2797
sp|P38110|TEL1_YEAST                MKAVQTKSNEERLKAY-LKITNEIKPQLRNFFFDSFP----DPLDWFEAKKTYTKGVAAS  2599
sp|Q13535|ATR_HUMAN                 -MLPKSAALSEKLKVFREFLLPRHPPIFHEWFLRTFP----DPTSWYSSRSAYCRSTAVM  2461
sp|P38111|MEC1_YEAST                -WQ--HTAVDGKLEFY-MEQVDKFPPILYQWFLENFP----DPINWFNARNTYARSYAVM  2211
PI4K_Plasmodium_falciparum_         --------------------------ISTIFN------IVFSDYIFEAKKNFIESHAAY  1401
tr|Q8I3V5|PI3K_Plasmodium_falciparum_ ---------------------------RQYFIDNSTCSSSPLGFDTEILQNFISSCAGY  1979
sp|Q8NEB9|PI3KC3_HUMAN              ---------------------------QNFFRKYAPSENGPNGISAEVMDTYVKSCAGY  732
                                                                                  .    :  . *

sp|Q13315|ATM_HUMAN                 SIVGYILGLGDRHVQNILINEQSAELVHIDLGVAFEQGKILPTPETVPFRLTRDIVDGMG  2859
sp|P38110|TEL1_YEAST                SIVGYILGLGDRHLNNILLDCSTGEPIHIDLGIAFDQGKLLPIPELVPFRLTRDIVDGFG  2661
sp|Q13535|ATR_HUMAN                 SMVGYILGLGDRHGENILFDSLTGECVHVDFNCLFNKGETFEVPEIVPFRLTHNMVNGMG  2524
sp|P38111|MEC1_YEAST                AMVGHILGLGDRHCENILLDIQTGKVLHVDFDCLFEKGKRLPVPEIVPFRLTPNLLDALG  2273
PI4K_Plasmodium_falciparum_         SLISYLLQVKDRHNGNLLLD-SDGHLIHIDYGFMLTNSPGNVNFETSPFKLTQEYLDIMD  1460
tr|Q8I3V5|PI3K_Plasmodium_falciparum_ SVITYILGIGDRHLDNLMVT-KDGRFFHIDFGYIFGEDPKPF---SPPMKLCKEMIEAMG  2035
sp|Q8NEB9|PI3KC3_HUMAN              CVITYILGVGDRHLDNLLLT-KTGKLFHIDFGYILGRDPKPL---PPPMKLNKEMVEGMG  788
                                    .:: ::* : ***  *::.    .. .*:* .  : ..         *::*  : :: :.

sp|Q13315|ATM_HUMAN                 ITGVEG--VFRRCCEKTMEVMRNSQETLLTIVEVLLYD-PLFDWTMNPLKALYLQQRPED  2916
sp|P38110|TEL1_YEAST                VTGVDG--LFRRSCERVYAVLRKDYVKVMCVLNILKWD-PLYSWVMSPVKKYEHLFEEEH  2718
sp|Q13535|ATR_HUMAN                 PMGTEG--LFRRACEVTMRLMRDQREPLMSVLKTFLHD-PLVEWSKPVK-----G--HSK  2574
sp|P38111|MEC1_YEAST                IIGTEG--TFKKSSEVTLALMRKNEVALMNVIETIMYD-RNMDHSIQKALKVLRN--KIR  2328
PI4K_Plasmodium_falciparum_         GEKSDNYEYFRRLIVSGFLEARKHSEEIILFVELMMPALKIPCFANGTQFCIESLKERFM  1520
tr|Q8I3V5|PI3K_Plasmodium_falciparum_ GAHSIGYEQFLKKCCLAYKYLRYHSQLIISLLDAMCDA-GLKDMK--------------  2079
sp|Q8NEB9|PI3KC3_HUMAN              GTQSEQYQEFRKQCYTAFLHLRRYSNLILNLFSLMVDA-NIPDIALEPDKTVKKVQDKFR  847
                                             * :         *     :: ... :

sp|Q13315|ATM_HUMAN                 ETELHPT----LN----ADDQECKRNLSDI----------------------          2951
sp|P38110|TEL1_YEAST                EITNFDNVSKFISNNDRNENQESYRALK------------------------          2746
sp|Q13535|ATR_HUMAN                 AP--LNETGEVVNEKAKTHVLDIEQRLQGVIK--------------------          2604
sp|P38111|MEC1_YEAST                GI--DPQDGLVLSVAGQTETLIQEAT--------------------------          2347
PI4K_Plasmodium_falciparum_         -----------TN----LTVDVCIQRINALIEASINNFRSVQYDYFQRITNGI          1558
tr|Q8I3V5|PI3K_Plasmodium_falciparum_ ----------------------------------------------------          2079
sp|Q8NEB9|PI3KC3_HUMAN              -----------LD----LSDEEAVHYMQSLIDESVHALF-------------          871
```

| | *Hs* ATM | *Sc* TEL1 | *Hs* ATR | *Sc* MEC1 | *Pf* PI4K | *Pf* PI3K | HsPI3K |
|---|---|---|---|---|---|---|---|
| 1: sp\|Q13315\|ATM_HUMAN | 100.0 | 43.1 | 34.6 | 31.4 | 21.2 | 24.4 | 24.6 |
| 2: sp\|P38110\|TEL1_YEAST | 43.1 | 100.0 | 33.7 | 32.0 | 28.1 | 27.9 | 26.0 |
| 3: sp\|Q13535\|ATR_HUMAN | 34.6 | 33.7 | 100.0 | 41.8 | 21.3 | 23.4 | 22.7 |
| 4: sp\|P38111\|MEC1_YEAST | 31.4 | 32.0 | 41.8 | 100.0 | 19.2 | 23.5 | 24.9 |
| 5: PI4K_Plasmodium_falciparum_ | 21.2 | 28.1 | 21.3 | 19.2 | 100.0 | 33.2 | 30.0 |
| 6: tr\|Q8I3V5\|PI3K_Plasmodium_falciparum_ | 24.4 | 27.9 | 23.4 | 23.5 | 33.2 | 100.0 | 49.3 |
| 7: sp\|Q8NEB9\|PI3KC3_HUMAN | 24.6 | 26.0 | 22.7 | 24.9 | 30.0 | 49.3 | 100.0 |

Figure S4:

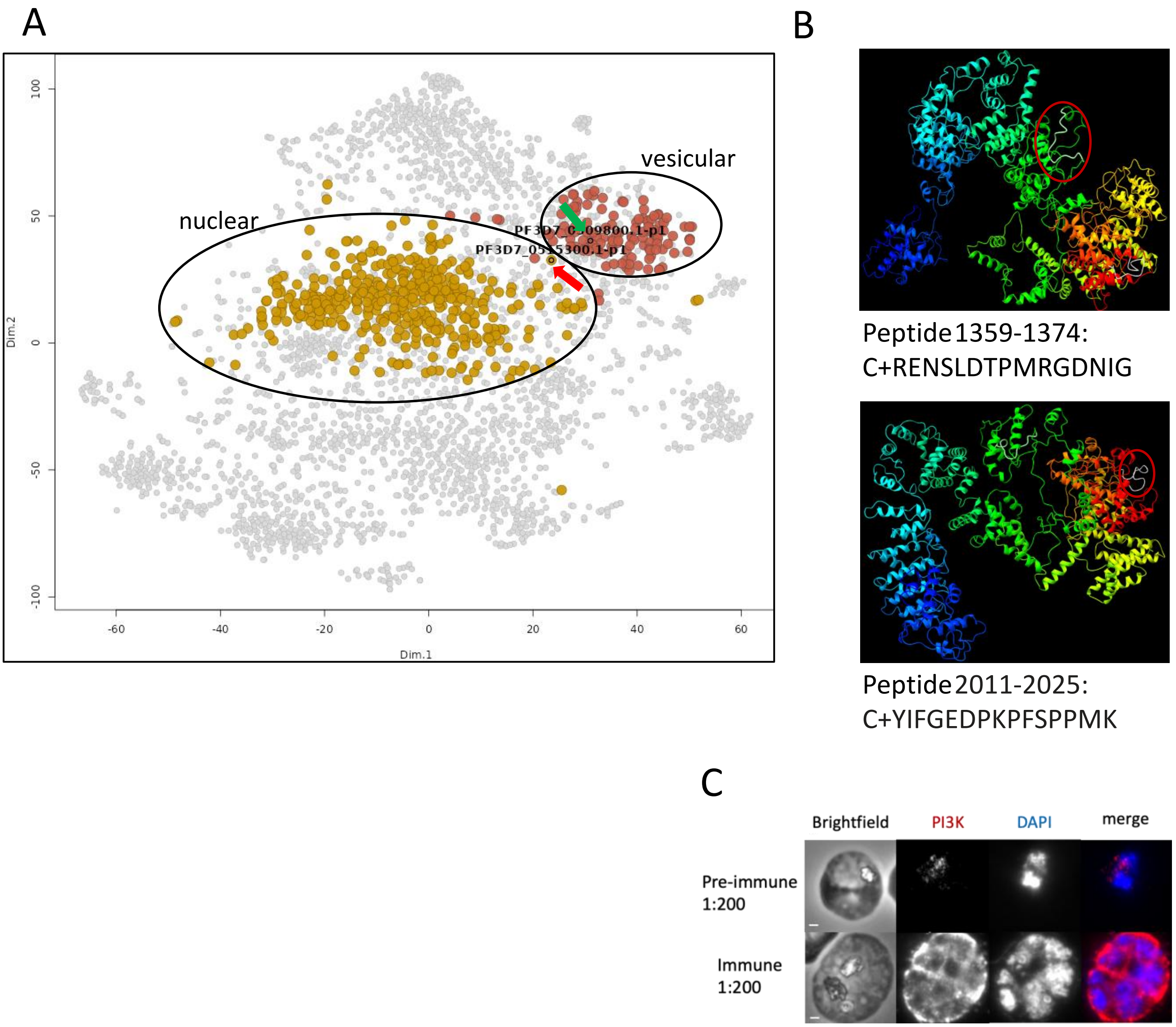
A
B
C
vesicular
nuclear
Dim.1
Dim.2
100
50
0
-50
-100
-60
-40
-20
0
20
40
60
Peptide 1359-1374:
C+RENSLDTPMRGDNIG
Peptide 2011-2025:
C+YIFGEDPKPFSPPMK
Brightfield
PI3K
DAPI
merge
Pre-immune
1:200
Immune
1:200

Figure S5A:

Control (iii) – all slices from Z-stack – scale bar 10um

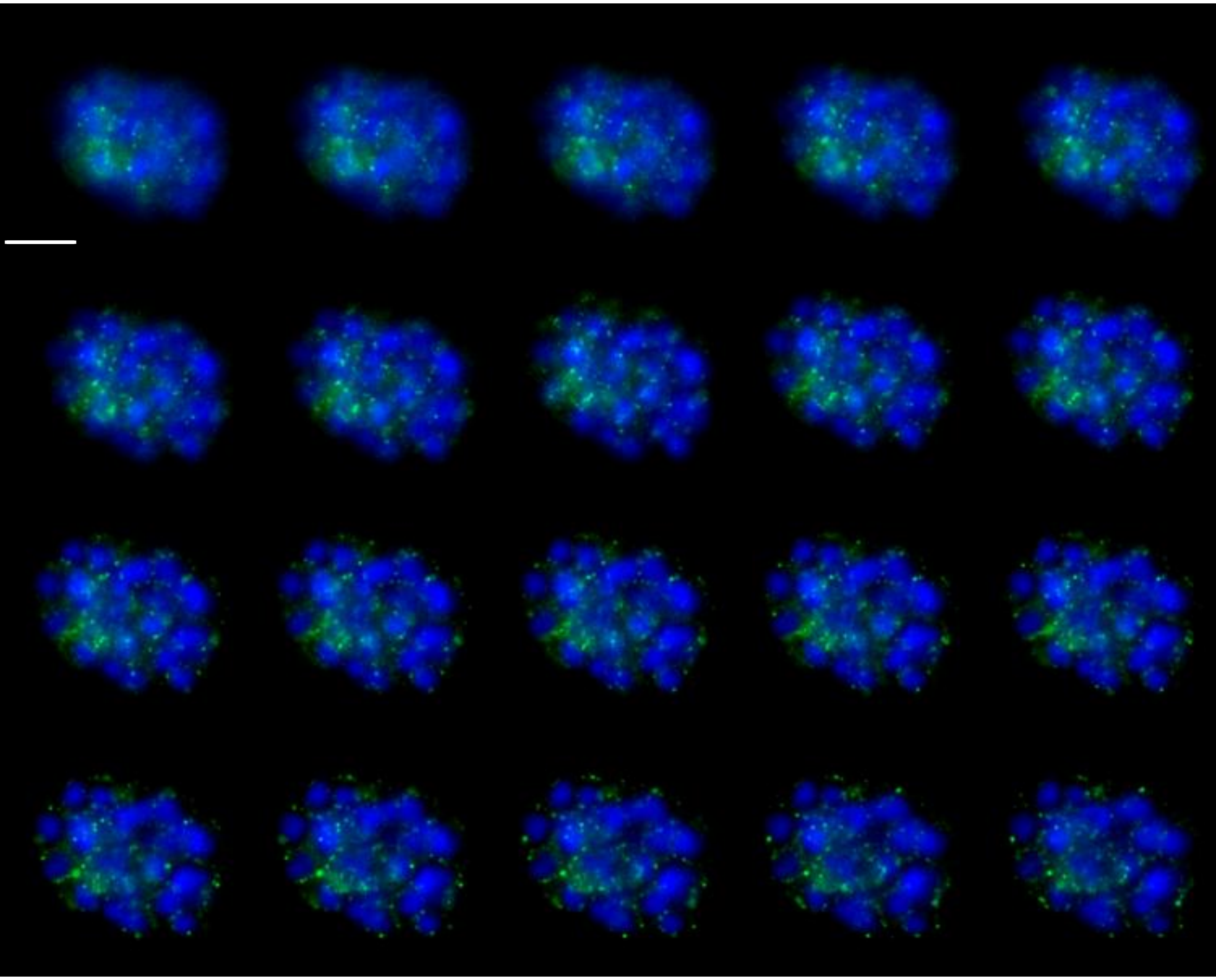

Figure S5B:

DHA (i) – all slices from Z-stack – scale bar 10um

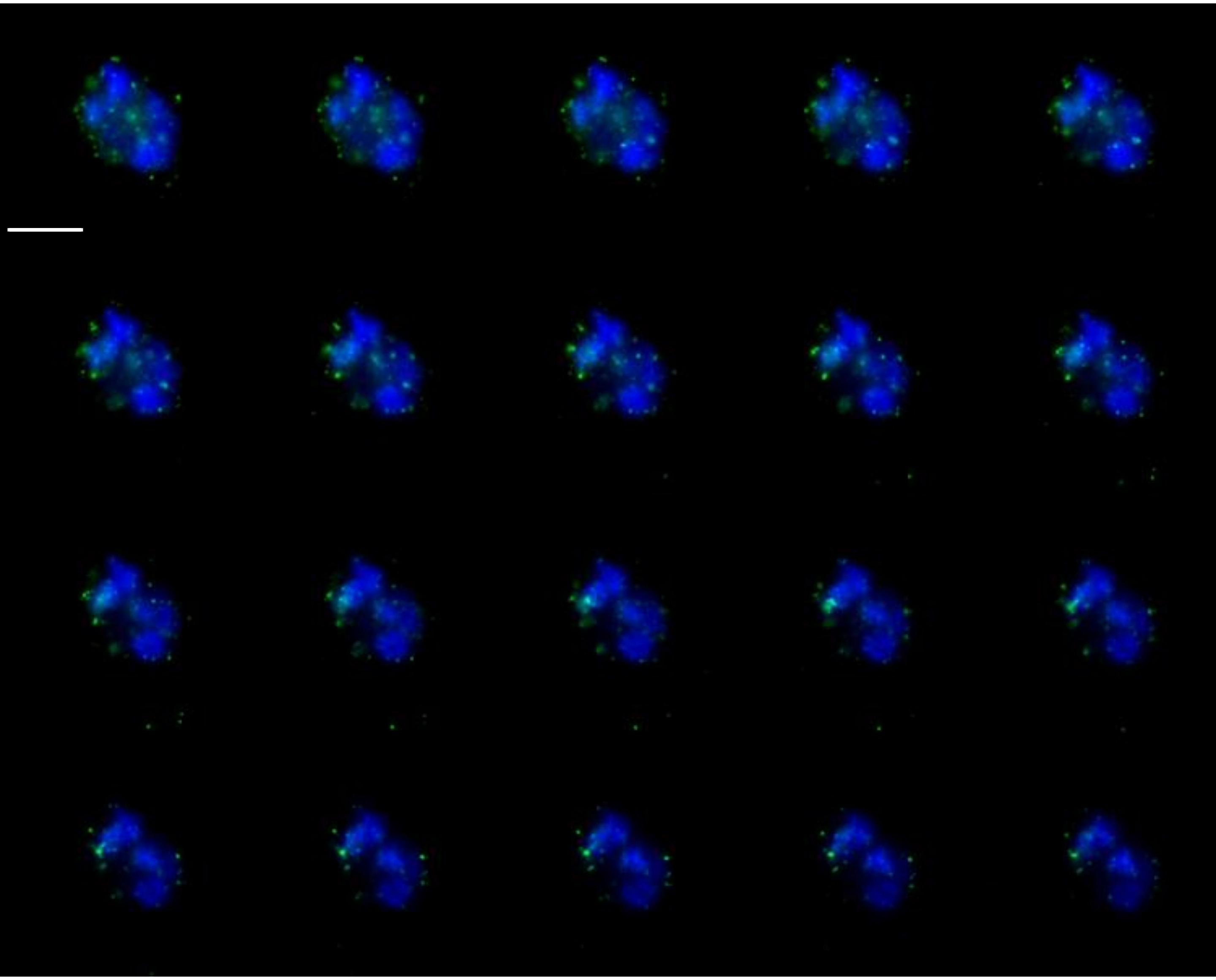

Figure S5C:

MMS (ii) – all slices from Z-stack – scale bar 10um

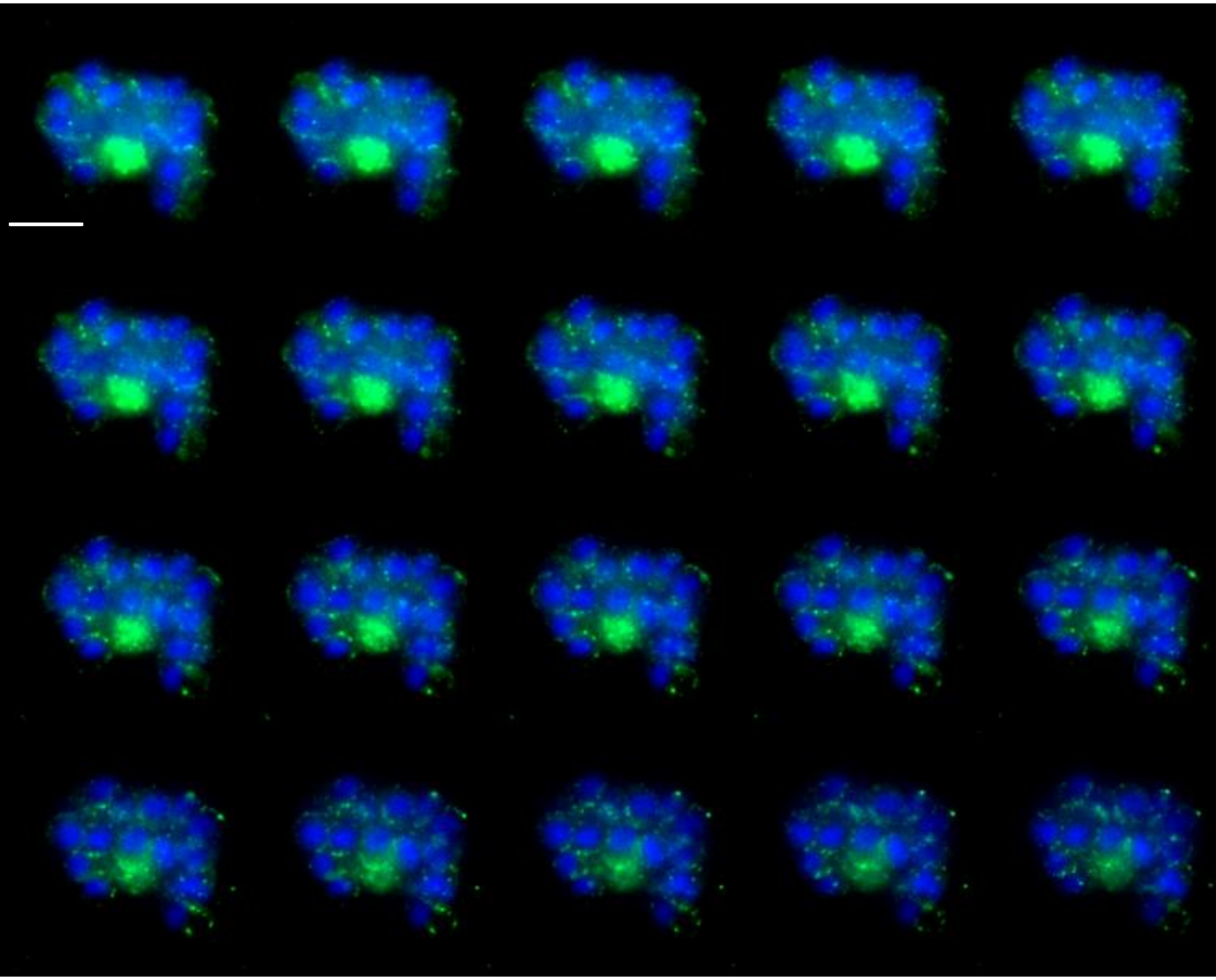

Figure S6:

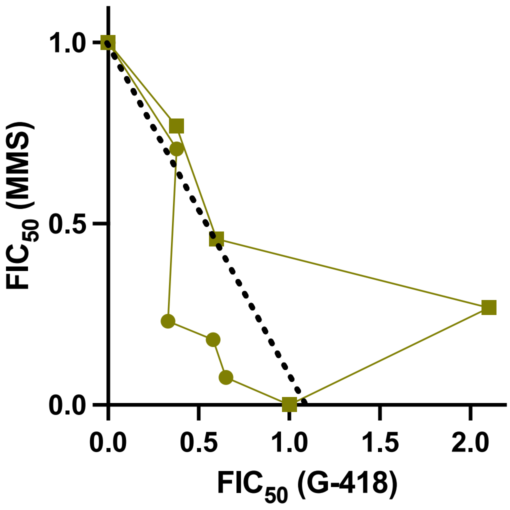


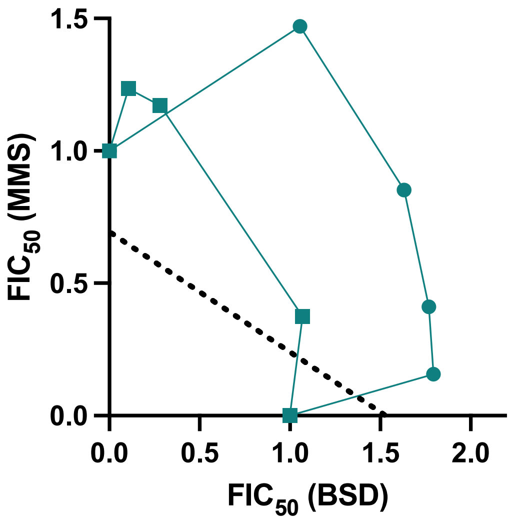


| MMS + drug | Sum of $FIC_{50}$ |
|---|---|
| Geneticin | 2.18(0.78-3.57) |
| Blasticidin | 1.85(1.41-2.28) |

Figure S7:

A

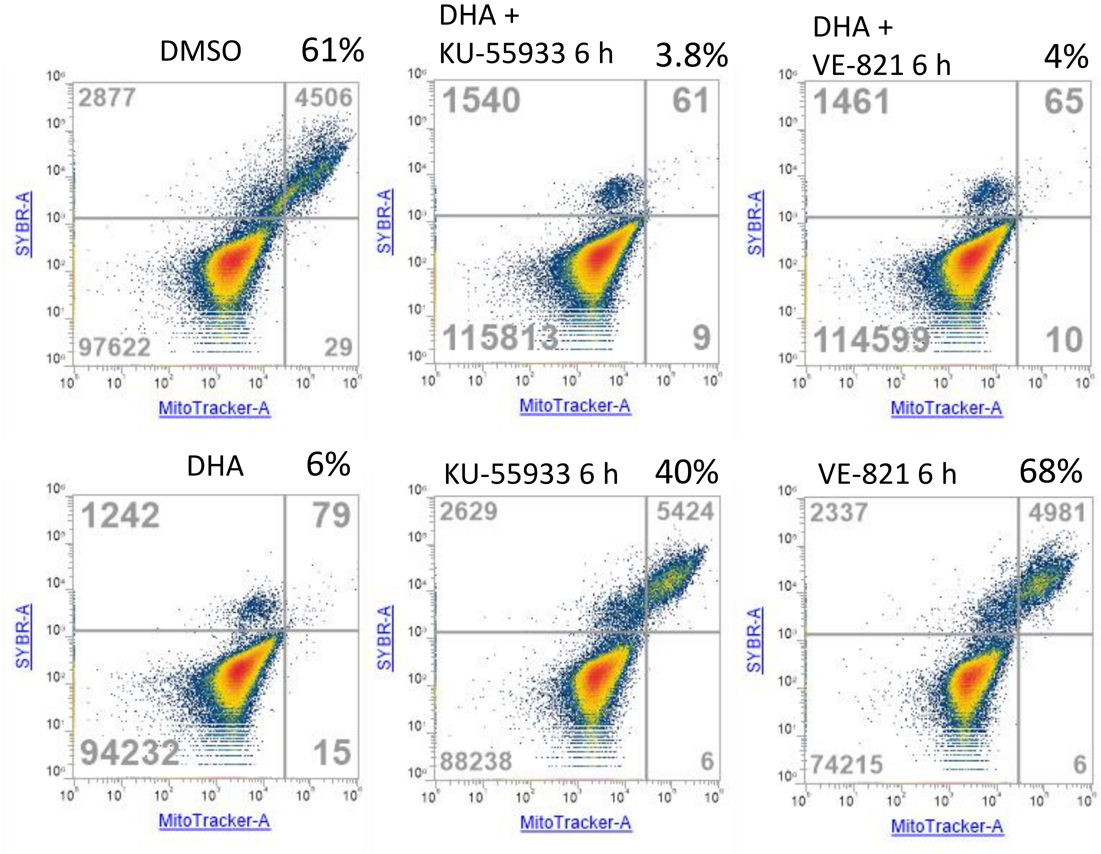


B

| | | Replicates $IC_{50}$ | | | | | | $IC_{50}$ mean | STDEV |
|---|---|---|---|---|---|---|---|---|---|
| MMS [µM] | 3D7 | 129.6 | 66.69 | 58.97 | 161.5 | 67.18 | | 97 | 46.0 |
| | MRA-1252 | 61.11 | 49.49 | 34.57 | 65.78 | 42.45 | 34.74 | 48 | 13.2 |
| | MRA-1240 | 173.8 | 124.1 | 302.6 | 66.09 | 71.19 | | 148 | 97.1 |
| VE-821 [nM] | 3D7 | 95.49 | 106.7 | 190.4 | | | | 131 | 51.9 |
| | MRA-1252 | 46.11 | 61.62 | 45.61 | | | | 51 | 9.1 |
| | MRA-1240 | 1065 | 1857 | 3139 | | | | 2020 | 1046.6 |
| KU-55933 [µM] | 3D7 | 44.25 | 32.31 | 35.1 | 49.77 | | | 40 | 6.2 |
| | MRA-1252 | 22.85 | 19.93 | | | | | 21 | 2.1 |
| | MRA-1240 | 31.43 | 19.38 | 26.16 | | | | 26 | 6.0 |

Figure S8A:

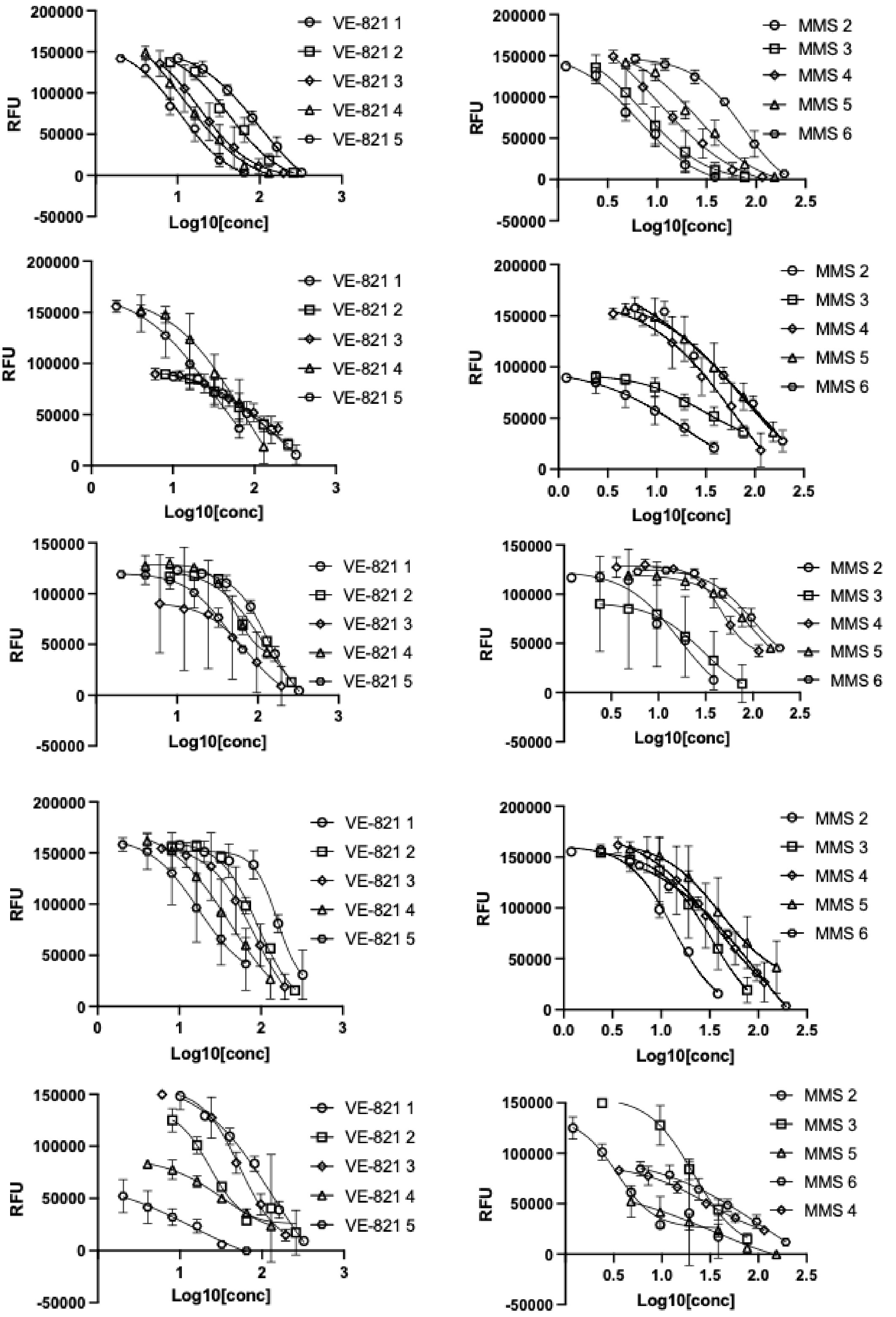
RFU
Log10[conc]
VE-821 1
VE-821 2
VE-821 3
VE-821 4
VE-821 5
RFU
Log10[conc]
MMS 2
MMS 3
MMS 4
MMS 5
MMS 6
RFU
Log10[conc]
VE-821 1
VE-821 2
VE-821 3
VE-821 4
VE-821 5
RFU
Log10[conc]
MMS 2
MMS 3
MMS 4
MMS 5
MMS 6
RFU
Log10[conc]
VE-821 1
VE-821 2
VE-821 3
VE-821 4
VE-821 5
RFU
Log10[conc]
MMS 2
MMS 3
MMS 4
MMS 5
MMS 6
RFU
Log10[conc]
VE-821 1
VE-821 2
VE-821 3
VE-821 4
VE-821 5
RFU
Log10[conc]
MMS 2
MMS 3
MMS 4
MMS 5
MMS 6
RFU
Log10[conc]
VE-821 1
VE-821 2
VE-821 3
VE-821 4
VE-821 5
RFU
Log10[conc]
MMS 2
MMS 3
MMS 5
MMS 6
MMS 4

Figure S8B:

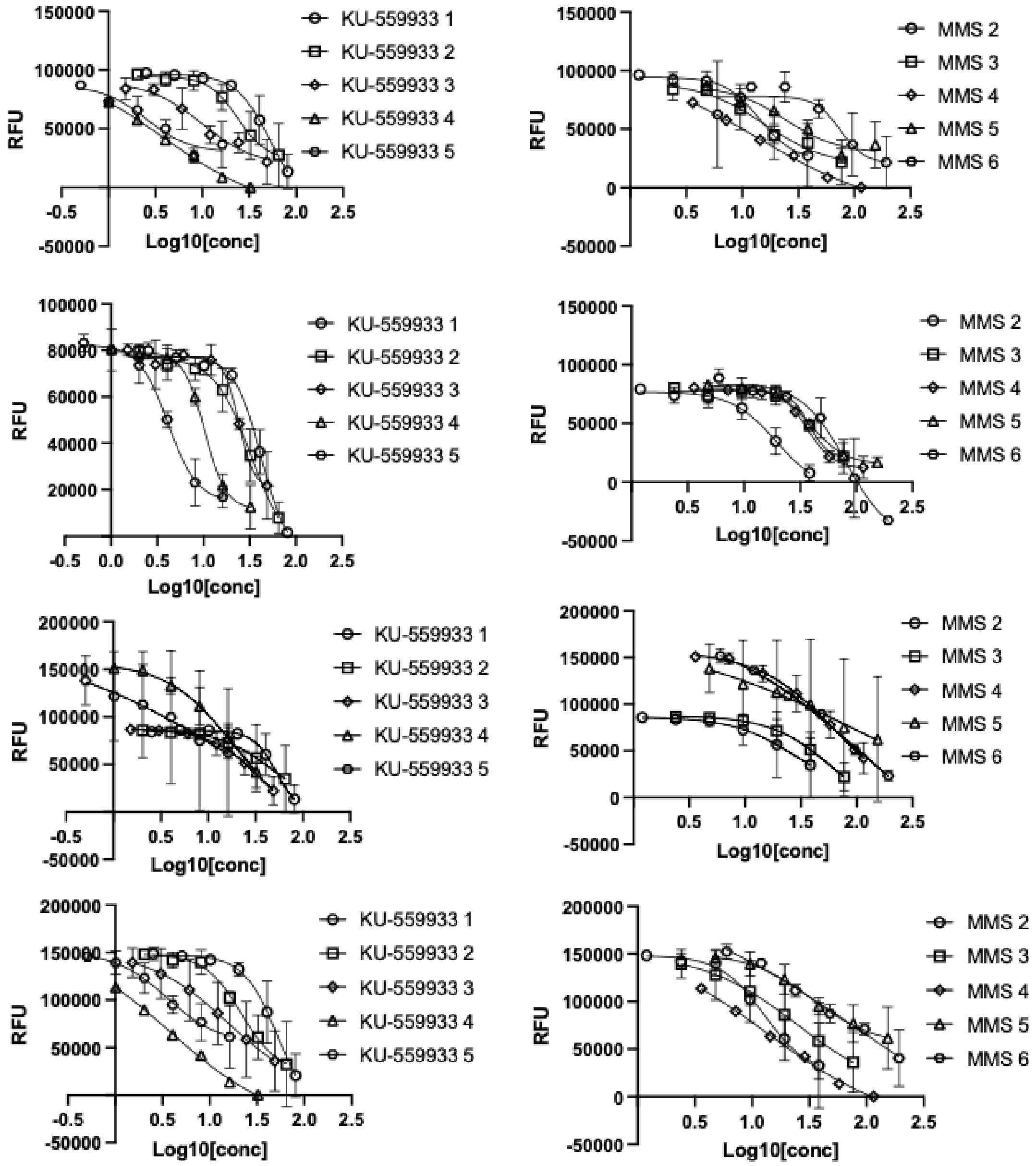
RFU
Log10[conc]
KU-559933 1
KU-559933 2
KU-559933 3
KU-559933 4
KU-559933 5
MMS 2
MMS 3
MMS 4
MMS 5
MMS 6

Figure S8C:

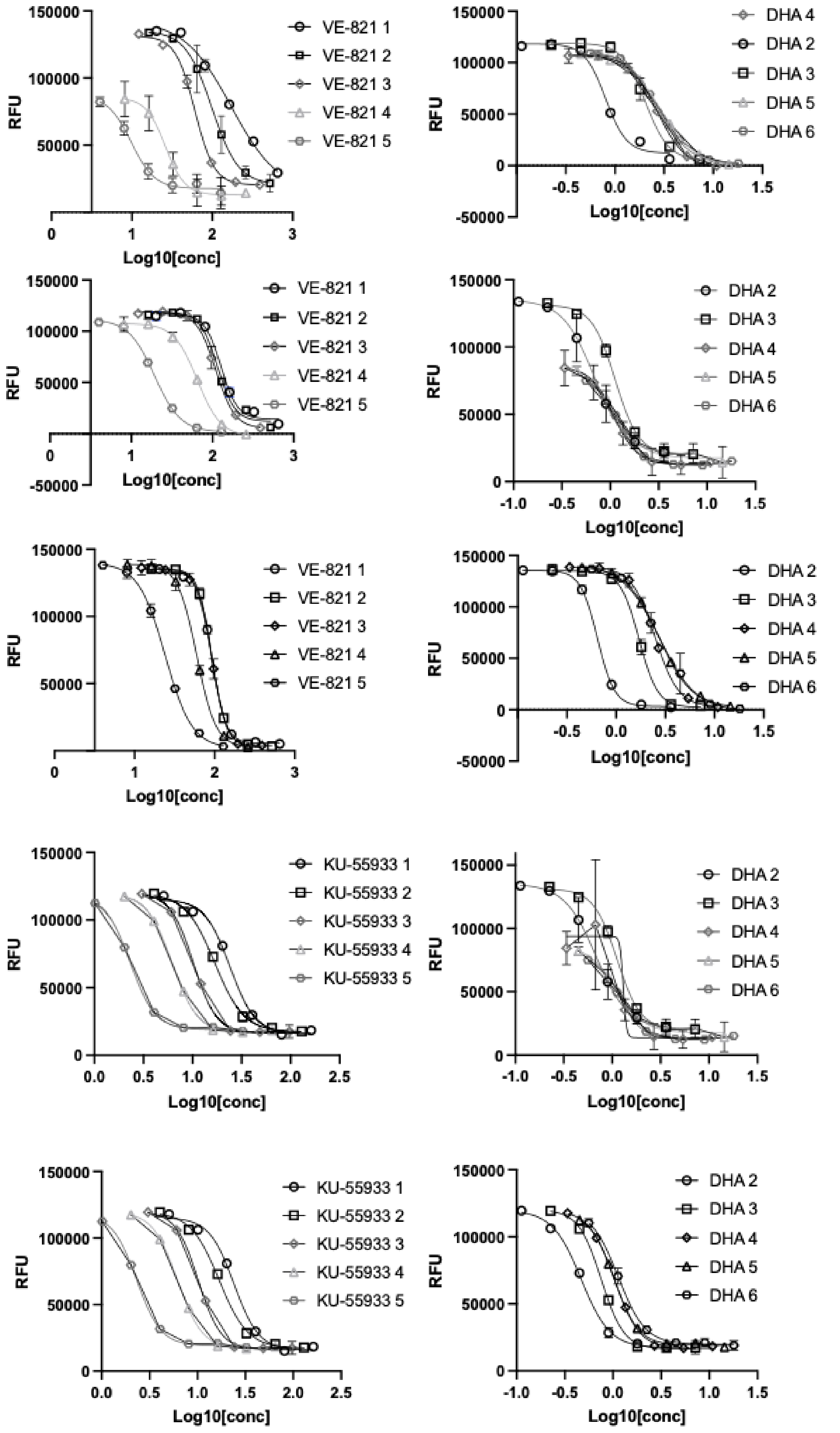
RFU
Log10[conc]
VE-821 1
VE-821 2
VE-821 3
VE-821 4
VE-821 5
DHA 4
DHA 2
DHA 3
DHA 5
DHA 6
RFU
Log10[conc]
VE-821 1
VE-821 2
VE-821 3
VE-821 4
VE-821 5
DHA 2
DHA 3
DHA 4
DHA 5
DHA 6
RFU
Log10[conc]
VE-821 1
VE-821 2
VE-821 3
VE-821 4
VE-821 5
DHA 2
DHA 3
DHA 4
DHA 5
DHA 6
RFU
Log10[conc]
KU-55933 1
KU-55933 2
KU-55933 3
KU-55933 4
KU-55933 5
DHA 2
DHA 3
DHA 4
DHA 5
DHA 6
RFU
Log10[conc]
KU-55933 1
KU-55933 2
KU-55933 3
KU-55933 4
KU-55933 5
DHA 2
DHA 3
DHA 4
DHA 5
DHA 6

Figure S8D:

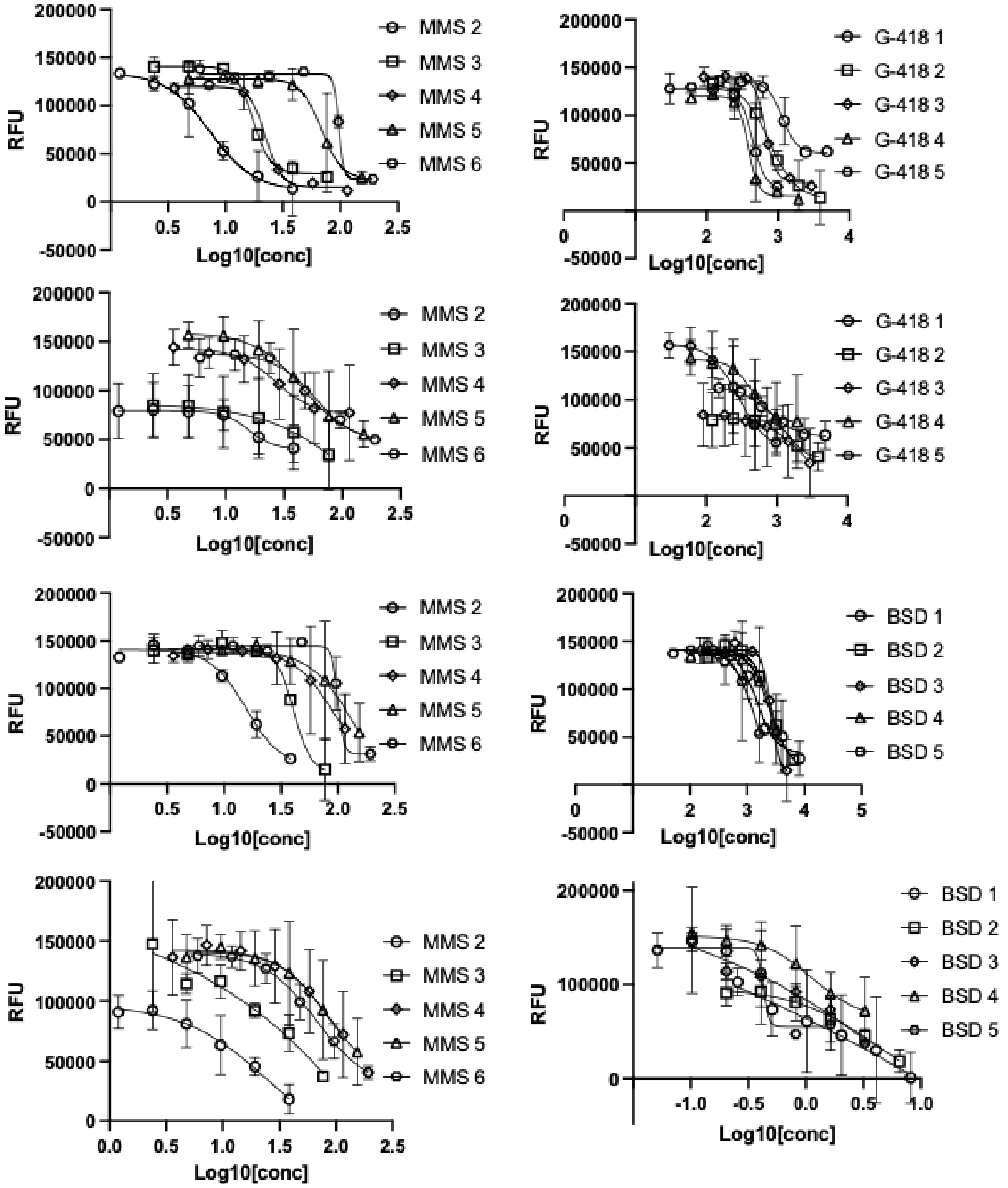
RFU
Log10[conc]
MMS 2
MMS 3
MMS 4
MMS 5
MMS 6
G-418 1
G-418 2
G-418 3
G-418 4
G-418 5
BSD 1
BSD 2
BSD 3
BSD 4
BSD 5
