## Supplementary figures and images for "Cell cycle checkpoint activity in the malaria parasite *Plasmodium falciparum*"

### Source

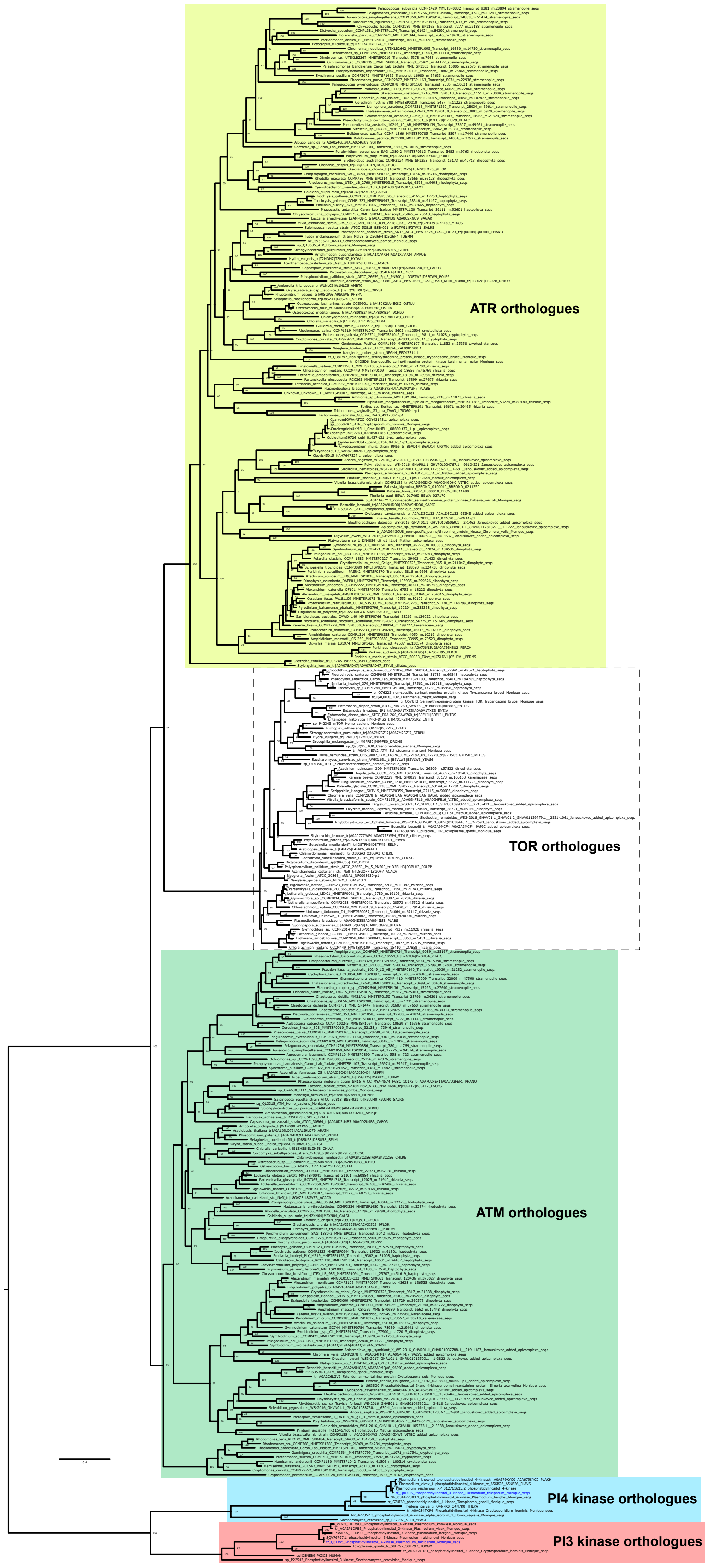
